# Saccades are coordinated with directed circuit dynamics and stable but distinct hippocampal patterns that promote memory formation

**DOI:** 10.1101/2022.08.18.504386

**Authors:** Isabella C. Wagner, Ole Jensen, Christian F. Doeller, Tobias Staudigl

## Abstract

During natural viewing, humans explore their environment by moving the eyes – a process that involves saccades to guide attention and shape memory. Saccades are controlled by visuo-oculomotor regions and are known to impact medial temporal lobe processing, but the circuit dynamics in humans are unclear. Here, we asked which of the functional routes between the visual cortex, frontal eye fields, and the hippocampus facilitates memory encoding during visual exploration and whether saccades affect neural representations. Forty-eight human participants underwent functional magnetic resonance imaging (fMRI) and continuous monitoring of eye gaze while studying scenes and performing a recognition memory test thereafter. We discovered saccade-related excitatory coupling from the visual cortex and frontal eye fields towards the hippocampus, independent of memory. This was complemented by hippocampal inhibition of the visual cortex that determined whether participants would later remember or forget. Moreover, saccades were associated with overall stable but distinct hippocampal voxel patterns during memory formation and the degree of pattern distinctiveness positively scaled with inhibition along the hippocampus-to-visual cortex route. This suggests that eye movements are timed to the orchestration of circuit connectivity and neural representations, setting the stage for memory formation.

## Introduction

Where we look, shapes what we remember. Humans and non-human primates actively gather information about their visual environment by moving the eyes. During free viewing, saccadic eye movements target distinct features of the visual scene to accumulate information (Henderson, 2017; Pertzov et al., 2009; Renninger et al., 2007) in support of memory (Bicanski and Burgess, 2019; Fehlmann et al., 2020; Lucas et al., 2019; Meister and Buffalo, 2016; Olsen et al., 2016). Direct electrophysiological recordings from both humans and non-human primates uncovered that saccades affect neuronal activity in the hippocampus (Doucet et al., 2020; Hoffman et al., 2013; Mao et al., 2021; Staudigl et al., 2022), surrounding medial temporal lobe regions (Ringo et al., 1994; Sobotka et al., 1997; Sobotka and Ringo, 1997), and the functional connectivity between them (Sobotka et al., 2002; Staudigl et al., 2022). Jutras and colleagues (2013) demonstrated that saccades induced a phase reset of hippocampal theta oscillations (3-12 Hz) when monkeys visually scanned images and that the reliability of this effect predicted subsequent recognition memory (Jutras et al., 2013). Moreover, saccades were shown to be phase-locked to alpha oscillations (7-14 Hz) in the occipital and medial temporal lobe as human participants studied visual scenes that were later remembered (Staudigl et al., 2017), and the frequency of visual sampling was associated with hippocampal activation levels recorded non-invasively using fMRI (Liu et al., 2017). These studies are in line with an active sensing account for vision where oculomotor actions and visual perception are tightly interlinked (Schroeder et al., 2010; Yang et al., 2016). Yet, human evidence as to whether saccades coincide with connectivity changes between visuo-oculomotor and hippocampal regions and thereby enhance memory formation is so far missing.

The relationship between visual exploration and memory appears to be bidirectional, reflecting two sides of the same coin: not only do we remember what we looked at, but what we remember also shapes where we look next (Kragel and Voss, 2022; Meister and Buffalo, 2016; Summerfield et al., 2006; Voss et al., 2017). Findings that substantiate this relationship revealed that eye movements depend on previous experience (Damiano and Walther, 2019) and that their efficiency relies upon the structural integrity of the hippocampus (Lucas et al., 2019; Olsen et al., 2016; Yoo et al., 2020). Recently, Kragel and colleagues discovered that phase-locking at the peak of hippocampal theta oscillations preceded eye movements towards remembered locations, indicating that hippocampal processing might modulate memory-guided viewing behavior (Kragel et al., 2020). Results from network modeling in non-human primates further suggested an effect of hippocampal activity on oculomotor regions (Ryan et al., 2020), building upon a distributed set of anatomical connections between the two systems (Shen et al., 2016). Together, this speaks for coordinated activity at the interface between visuo-oculomotor and medial temporal regions but it is unclear whether such bidirectional interactions support human memory.

In this experiment, 48 human participants studied scene photographs while undergoing functional magnetic resonance imaging (fMRI) and continuous eye tracking. Following the study period, participants were tested for their recognition memory by dissociating old from novel scenes (**Figure 1A**). Our main focus was the saccade-related and directed connectivity within a network of brain regions known to be involved in visual processing, oculomotor control (Johnston and Everling, 2008; Munoz and Everling, 2004) and memory formation (Kim, 2011; Spaniol et al., 2009), including the visual cortex (VC), the frontal eye fields (FEF) and the hippocampus (HC). We hypothesized that network activity would be initiated (and thus be “driven”) by saccades, either via oculomotor activity expressed within FEF, via visual input arriving in VC, or through a combination of both. Connectivity between visuo-oculomotor regions and the hippocampus was expected to be bidirectional, in line with previous work showing reciprocal links between visual exploration and memory (Kragel and Voss, 2022; Meister and Buffalo, 2016; Voss et al., 2017), and the strength of these connections should depend on participants’ trial-by-trial performance in the subsequent memory test. To scrutinize these predictions, we developed a saccade-related analysis approach that leveraged our fast fMRI acquisition sequence and employed dynamic causal modeling (DCM; Stephan and Friston, 2010).

**Figure 1.**
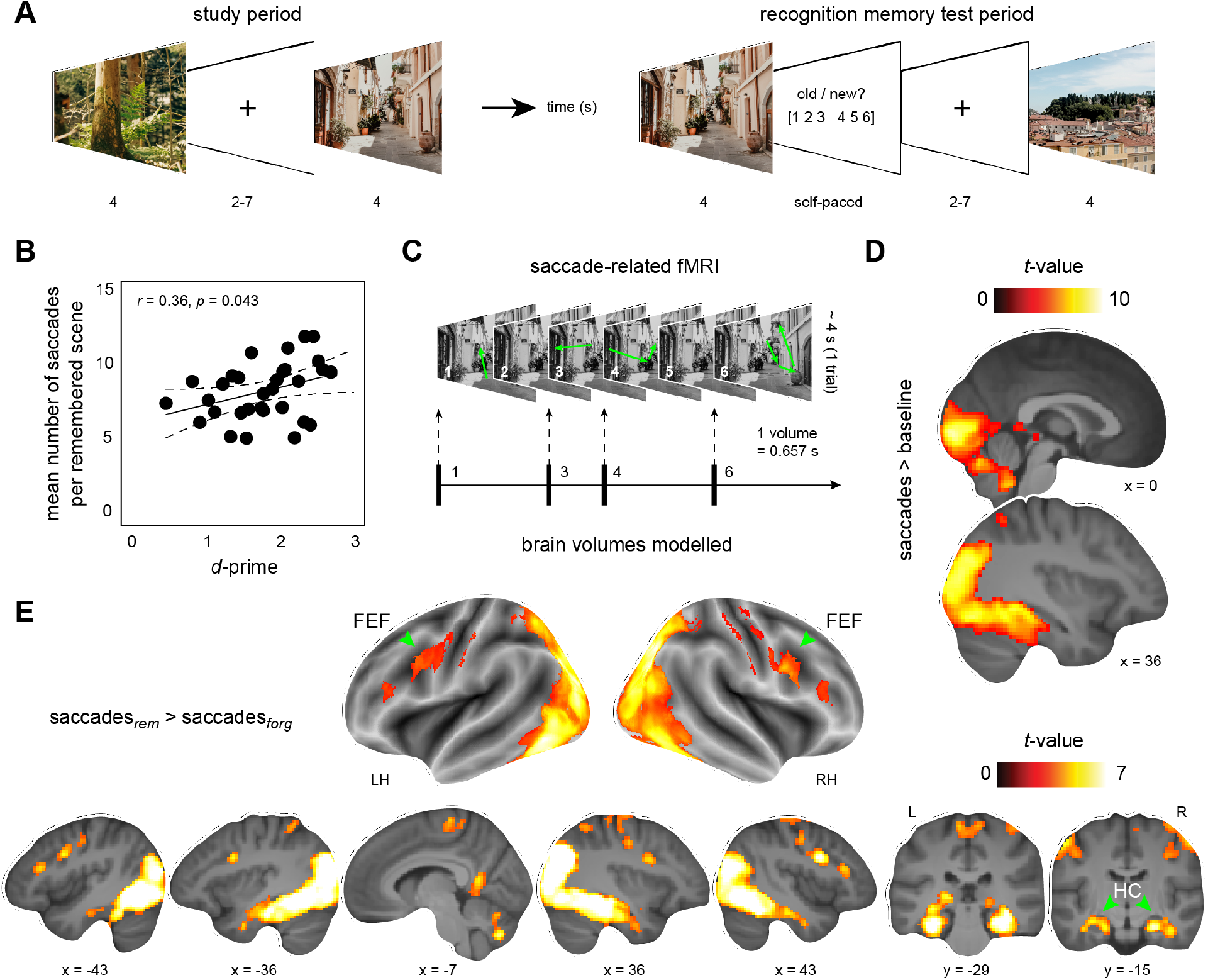
Recognition memory task and saccade-related fMRI. **(A)** Participants studied colored scene photographs and completed a recognition memory test thereafter (panels **A** and **C**: note that due to copyright reasons actual stimulus material is replaced by scenes that are openly available on Unsplash, photographed by Liza Rusalskaya, https://unsplash.com). **(B)** Positive cross-participant correlation of the mean number of saccades per remembered scene and individual recognition memory performance (*d*-prime). **(C)** Saccade-related fMRI approach: to capture the effect of saccadic eye movements on memory formation (example saccade trajectories indicated in green), we explicitly modelled brain volumes during which saccades were produced (e.g., volumes 1, 3, 4, 6; **Materials and Methods**). **(D)** Brain activation profile demonstrating the general effect of saccades compared to the implicit fixation baseline (contrast [saccades_rem_ ∩ saccades_forg>_] > baseline, **Table S1, Figure S1**), and **(E)** during memory formation (**Table S2**). All results are shown at *p* < 0.05 FWE-corrected at cluster level (cluster-defining threshold of *p* < 0.001). Brain slice images are based on the average structural scan of all participants. FEF, frontal eye fields, HC, hippocampus, LH, left hemisphere, RH, right hemisphere, L, left, R, right.

Furthermore, eye movements might reflect overt attention towards specific features of a visual scene. Previous work has shown that attention modulates the similarity (or “stability”) of distributed HC representations (or, activity in voxel patterns) and promotes memory (Aly and Turk-Browne, 2016, 2015). However, this work left unanswered whether the stabilizing effect of attention could actually be tied to saccade-related HC activity. To investigate this, we performed representational similarity analysis (RSA; Kriegeskorte et al., 2008; Wagner et al., 2016) and quantified the degree to which saccades were related to changes in HC pattern similarity as participants studied scenes that were subsequently remembered or forgotten. By reconciling results from network modelling and multivariate pattern analysis, we provide evidence that saccade-related HC patterns are stable but distinct in the service of memory formation, and that the distinctiveness of HC patterns specifically scales with directed connectivity from the HC towards the visual system.

## Results

### Recognition memory performance is higher when participants make more saccades

Following the initial study period where participants encoded scene photographs, they were asked to complete a recognition memory test during which they dissociated old from novel scenes (i.e., 200 old scenes intermixed with 100 new scenes; **Figure 1A**). Participants remembered more than two thirds of the material correctly [∼76%, mean ± SEM, 226.7 ± 5.4] and produced on average 7.79 (± 0.34) saccades per trial. The number of saccades during later remembered scenes (8.38 ± 0.34) was significantly higher compared to later forgotten scenes (7.2 ± 0.36; paired-samples *t*-test, *N* = 32; *t*(31) = 8.101, Cohen’s *d* = 1.43, 95% CI = [0.88, 1.47], *p*_*two-tailed*_ < 0.0001). This was also reflected in a positive cross-participant relationship between individual recognition performance (*d*-prime, 1.82 ± 0.12) and the average number of saccades when studying later remembered scenes (*r*_*Pearson*_ = 0.36, 95% confidence interval (CI) = [0.01, 0.63], *p*_*two-tailed*_ = 0.043; **Figure 1B**). In other words, participants who made more saccades to visually explore the encoded scenes were better able to dissociate old from novel material during the subsequent recognition memory test.

### Saccades are associated with activation changes that support memory formation

We next turned to the fMRI data and tested for saccade-related changes in brain activation during later remembered compared to later forgotten trials. Specifically, we reasoned that saccades would be associated with activation increases in brain regions implicated in visual processing, oculomotor control and memory including the occipital lobe, the frontal eye fields (FEF), the hippocampus (HC) and surrounding medial temporal lobe regions (Johnston and Everling, 2008; Kim, 2011; Liu et al., 2017; Munoz and Everling, 2004; Spaniol et al., 2009). To test this, we developed a saccade-related approach that leveraged our fast MRI acquisition sequence and that allowed us to identify time points and associated brain images (i.e., volumes) during which saccades took place (**Figure 1C, Materials and Methods**). Saccade-related volumes were then grouped into such that occurred during subsequently remembered or forgotten scenes (saccades_rem_, saccades_forg_) and were contrasted to test for activation increases associated with successful memory formation.

As expected, saccades were linked to bilateral activation increases in occipital, inferior temporal and parietal regions (one-sample *t*-test, *N* = 32, contrast [saccades_rem_ ∩ saccades_forg_] > baseline; **Figure 1D**), extending into the parahippocampal gyrus, the lateral geniculate region, and the lateral prefrontal cortex (**Figure S1, Table S1**). We next focused on saccade-related processes during successful memory formation. Saccades during encoding of later remembered (compared to later forgotten) scenes were associated with an activation profile similar to the above but additionally included the HC and retrosplenial cortex, the FEF, and adjacent premotor and somatosensory regions (one-sample *t*-test, *N* = 32, contrast saccades_rem_ > saccades_forg_; **Figure 1E, Table S1**).

To validate that the saccade-related approach provided a better model fit to our data than other, more conventional models, we also analyzed the data by parametrically modulating trial-based activation changes with the number of saccades that occurred during each scene presentation (parametric model; **Figure S2, Table S2, Materials and Methods**). Model comparison using Bayesian Model Selection (BMS) revealed that the saccade-related model provided a better model fit compared to the parametric model in ∼90% of all voxels in the brain (**Figure S3**), including a higher average model fit within the VC, the FEF, and the HC [**Results S1**; and see **Figure S2** and **Table S3** for analysis with a conventional model that was oblivious to saccades but only took entire trials (i.e., visual scene presentations) into account].

### Saccades drive network activity via visual cortex and frontal eye fields

Our central question was whether saccades were related to directed connectivity from visuo-oculomotor regions towards the HC (and vice versa, given the bidirectional link between visual exploration and memory), and whether connectivity would be associated with memory formation. We hence focused on interactions within a VC-FEF-HC network and performed dynamic causal modeling (DCM; Stephan and Friston, 2010). We defined regions-of-interest (ROIs) based on the abovementioned findings of saccade-related activation changes (**Materials and Methods**). To reduce model space complexity (Stephan et al., 2010), we grouped ROIs into left- and right-lateralized networks, initially performed DCM analysis in the left hemisphere (**Figure 2A**) and subsequently validated results in the right hemisphere. To briefly explain, we composed a basic network model (**Figure 2B**) that consisted of three nodes that were fully connected (i.e., bilateral intrinsic connections between all ROIs and self-connections) and that systematically varied depending on which of the intrinsic between-ROI connections were modulated by subsequent memory performance (i.e., effect of MEMORY, saccades_rem_ > saccades_forg_). We hypothesized that network activity would be initiated (and thus be “driven”) by saccades, either via visual input arriving in VC (family I), via oculomotor activity expressed within the FEF (family II), or through a combination of both (family III; effect of SACCADE; **Figure 2C**). This resulted in 3 × 64 models that were grouped into three model families with respect to the exact location of the driving input.

**Figure 2:**
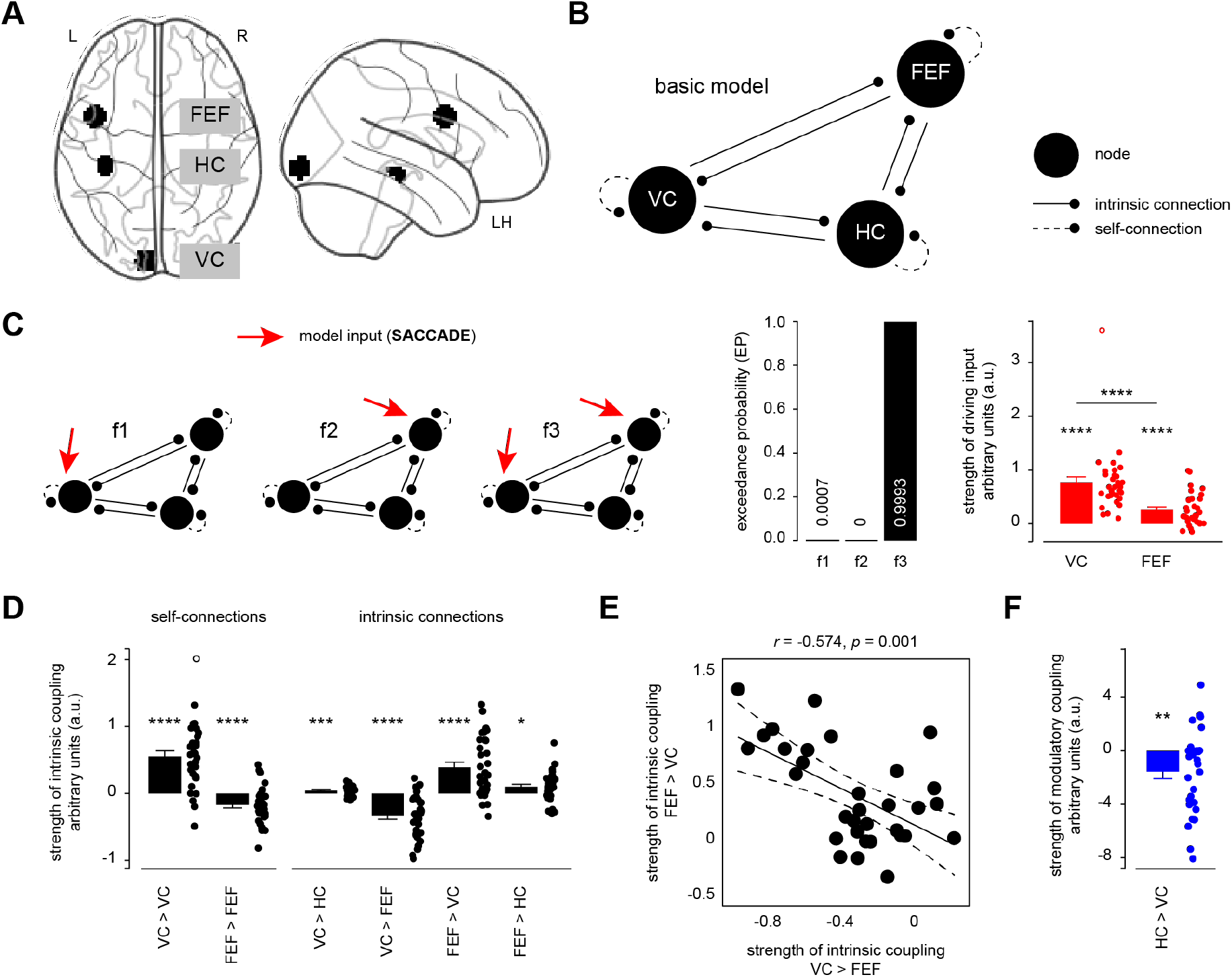
Directed connectivity within a VC-FEF-HC network is coordinated with saccades and promotes memory formation. **(A)** Left-lateralized network of regions-of-interest (ROIs) that included the visual cortex (VC), frontal eye fields (FEF), and the hippocampus (HC). **(B)** Basic model including bilateral intrinsic connections between all nodes that could be modulated by MEMORY (saccades_rem_ > saccades_forg_), and self-connections (these could not be modulated). Randomization of the number of modulatory connections yielded a basic model space with 64 models (not shown). **(C)** Network activity (SACCADE; saccade > baseline) could enter the model vial visual input arriving in VC, oculomotor activity expressed within FEF, or both. Models were grouped into three model families (f1-f3) and Bayesian Model Selection (BMS) revealed that f3 best explained the data. **(D)** Strength of intrinsic coupling parameters for self- and intrinsic connections. **(E)** Negative cross-participant correlation between coupling strength of the FEF-to-VC and VC-to-FEF connections. **(F)** Coupling strength of the inhibitory HC-to-VC connection that was modulated by MEMORY (saccades_rem_ > saccades_forg_). Error bars show SEM and data points reflect individual participant values. Hollow data points were formally identified as outliers; results remained robust after excluding these and repeating the analyses. LH, left hemisphere, **** *p* < 0.0001, *** *p* < 0.001, ** *p* < 0.01, * *p* < 0.05.

Bayesian Model Selection (BMS) showed that family III was most likely associated with the data [exceedance probability (EP): family I, 0.0007; family II, 0; family III, 0.9993; **Figure 2C**]. Put differently, the network was driven by saccade-related activity in both the VC [strength of driving input indexed through arbitrary units (a.u.), mean ± SEM: 0.77 ± 0.11; one-sample *t*-test, *N* = 32; *t*(31) = 7.299, Cohen’s *d* = 1.29, 95% CI = [0.55, 0.98], *p*_*two-tailed*_ < 0.0001] and FEF [0.26 ± 0.05; one-sample *t*-test, *N* = 32, *t*(31) = 4.987, *d* = 0.88, 95% CI = [0.15, 0.37], *p*_*two-tailed*_ < 0.0001]. Comparing VC and FEF directly, we found stronger VC than FEF driving input [paired-sample *t*-test, *N* = 32; *t*(31) = 4.596, *d* = 0.81, 95% CI = [0.28, 0.73], *p*_*two-tailed*_ < 0.0001; **Figure 2C**].

### Visual cortex initiates saccade-related activity in the hippocampus

Having established that saccades drove network activity via the VC and FEF, we went on to investigate the directed connectivity within the VC-FEF-HC network. Our aim was to assess the specific network structure as indexed by intrinsic functional connectivity (i.e., changes in between-ROI connectivity related to saccades). Apart from our hypothesis that saccades would increase connectivity from the VC and/or the FEF towards the HC, we were agnostic to the specific network (or model) structure and applied Bayesian Model Averaging (BMA) across the 64 models that comprised family III (**Materials and Methods**). This procedure yielded average model parameters that were weighted by the posterior probability of each of the models in the model space.

Results demonstrated a significant excitatory connection from the VC to the HC [strength of intrinsic coupling measured in arbitrary units (a.u.), mean ± SEM: 0.04 ± 0.01; one-sample *t*-test, *N* = 32; *t*(31) = 3.79, *d* = 0.67, 95% CI = [0.02, 0.07], *p*_*two-tailed*_ = 0.0007, α_Bonferroni_ = 0.05/9 = 0.005; **Figure 2D**], showing that the VC increased activity in the HC when saccades occurred. Also, the FEF showed an excitatory connection towards the hippocampus [0.1 ± 0.04; one-sample *t*-test, *N* = 32; *t*(31) = 2.607, *d* = 0.46, 95% CI = [0.21, 0.18], *p*_*two-tailed*_ = 0.014; but note that this latter result did not survive Bonferroni-correction].

Findings further revealed a bilateral connection between the VC and the FEF: the VC inhibited the FEF [-0.33 ± t0.06; one-sample *t*-test, *N* = 32; *t*(31) = -5.924, *d* = -1.05, 95% CI = [-0.45, -0.22], *p*_*two-tailed*_ < 0.0001; **Figure 2D**], while the latter showed an excitatory connection towards the VC [0.39 ± 0.08; one-sample *t*-test, *N* = 32; *t*(31) = 5.032, *d* = 0.89, 95% CI = [0.23, 0.55], *p*_*two-tailed*_ < 0.0001; **Figure 2D**]. These intrinsic coupling parameters were negatively correlated (*r*_Pearson_ = -0.574, 95% CI = [-0.77, -0.28], *p*_*two-tailed*_ = 0.001; **Figure 2E**) such that stronger saccade-related inhibitory VC-to-FEF coupling was associated with stronger excitatory FEF-to-VC coupling.

### Saccade-related inhibition of visual cortex by the hippocampus during memory formation

The major goal of our analysis was to probe whether saccades enhanced memory formation by affecting connectivity within the VC-FEF-HC network. We tested this by focusing on the modulatory connections (MEMORY, saccades_rem_ > saccades_forg_) that we obtained from BMA (see above).

Results showed significant inhibitory coupling that originated from the HC and targeted the VC [strength of modulatory coupling measured in arbitrary units (a.u.), mean ± SEM: -1.55 ± 0.53; one-sample *t*-test, *N* = 32; *t*(31) = -2.913, *d* = -0.51, 95% CI = [-2.64, -0.47], *p*_*two-tailed*_ = 0.006; α_Bonferroni_ = 0.05/6 = 0.008; **Figure 2F, Figure 3**]. Said differently, activity in the HC decreased VC activity when participants visually encoded scenes that were later remembered (compared to when participants visually encoded scenes that were later forgotten). None of the other connections were significantly modulated by MEMORY.

**Figure 3:**
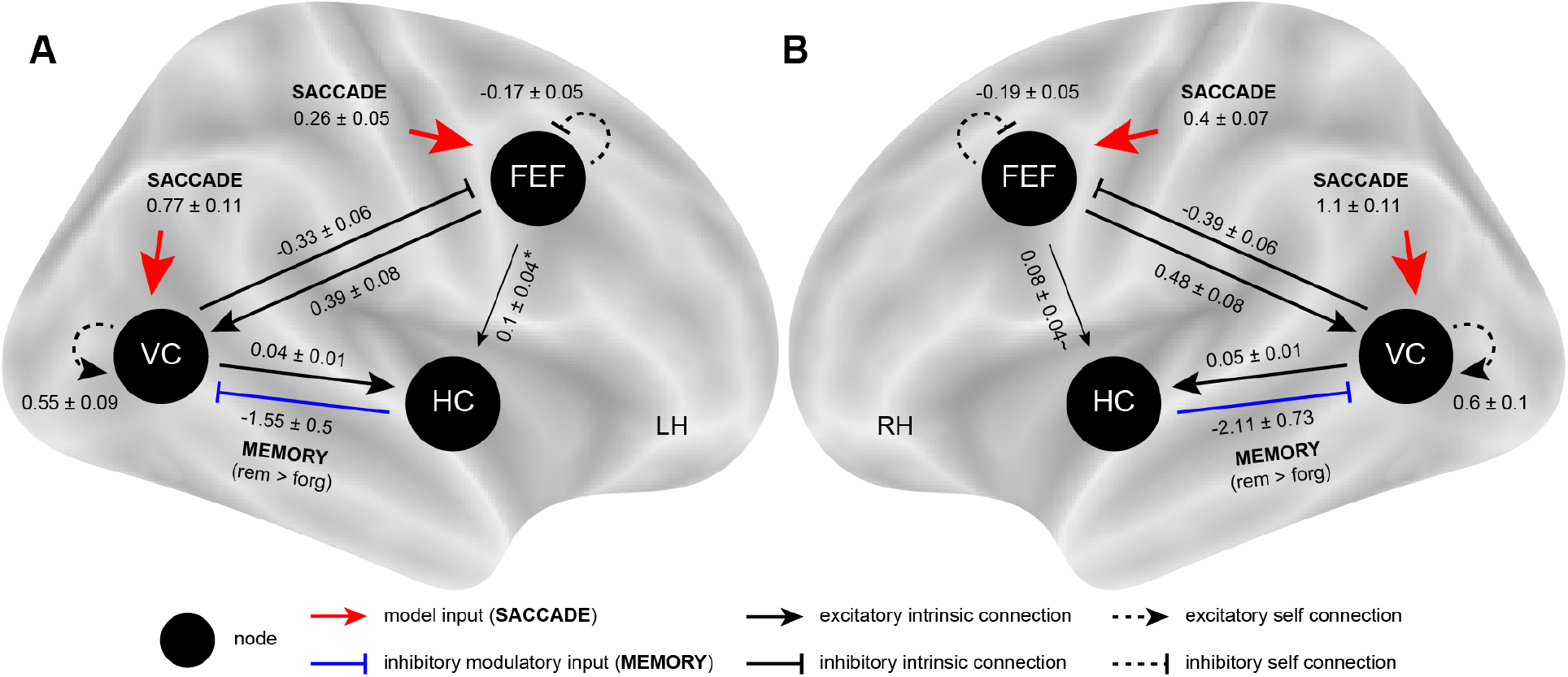
Final DCM network structure after Bayesian Model Averaging. **(A)** We initially performed DCM in a left-lateralized network that included the visual cortex (VC), frontal eye fields (FEF), and the hippocampus (HC). Bayesian Model Averaging (BMA) was used to obtain average model parameters that were weighted by the posterior probability of each of the 64 models within family III (input of SACCADE into both the VC and FEF). **(B)** Results were fully replicated in the right-lateralized network. **(A-B)** The FEF-HC intrinsic connection (indicated with thinner arrow) did not survive correction for multiple comparisons in the left hemisphere (LH, indicated through *) and showed a tendency towards significance (*p* = 0.078) in the right hemisphere (RH, indicated through ∼). Coupling strength (arbitrary units, a.u.) is represented as mean ± SEM.

### Control analyses and whole-brain connectivity

To validate the DCM results, we repeated our analyses with the same (but mirrored) ROIs in the right hemisphere (all other parameters were identical to the main analysis, see also **Materials and Methods**). Corroborating our left-lateralized findings above, BMS on right-lateralized data revealed that family III (input into both the VC and FEF) explained the data best [exceedance probability (EP): family I, 0.0002; family II, 0; family III, 0.9998]. Using the remaining model space of 64 models, we performed BMA to obtain the average model parameters. Results revealed a virtually identical network structure in the right hemisphere (**Figure 3B, Table S4**).

Moreover, to uncover the whole-brain connectivity profiles of the VC, FEF, and HC during saccade-related memory formation rather than just focusing on their local connectivity with the confined DCM networks, we also performed psychophysiological interaction analysis (PPI; Friston et al., 1997). Results showed wide-spread connectivity changes during saccade-related memory formation for all of the three seed regions (**Figure S4, Table S5**)

### Stable but distinct saccade-related patterns in the hippocampus and frontal eye fields during memory formation

So far, we have demonstrated that saccades affected the direction of information flow within a VC-FEF-HC network during memory formation. Moving forward, we drew upon work that previously demonstrated the “stabilizing” effect of attention on neural patterns in the HC and its beneficial impact on memory (Aly and Turk-Browne, 2016, 2015). What remained unclear, however, is whether such increases in pattern stability (or similarity) could be attributed to the influence of saccades on HC activity, since saccades may reflect allocation of overt attention. Our reasoning was two-fold (see also matrix in **Figure 4**): (1) Activation patterns linked to saccades during later remembered scenes might be more stable (or more similar to each other, indicating consistent processing or a high within-condition pattern similarity) than saccade-related patterns during later forgotten scenes (lower within-condition pattern similarity, contrast W_rem_ > W_forg_). (2) Additionally, saccade-related patterns during later remembered scenes might be more distinct from later forgotten material (or dissimilar, indicating differential processing or a higher within-than between-condition similarity, contrast W_rem_ > B_rem*forg_), which could facilitate subsequent recognition (LaRocque et al., 2013). To tackle this, we performed whole-brain, searchlight-based representational similarity analysis (RSA; Kriegeskorte et al., 2008; Wagner et al., 2016) and quantified the degree to which local voxel patterns were stabilized and/or distinct as participants produced saccades to study visual scenes that were subsequently remembered or forgotten (**Materials and Methods**).

**Figure 4.**
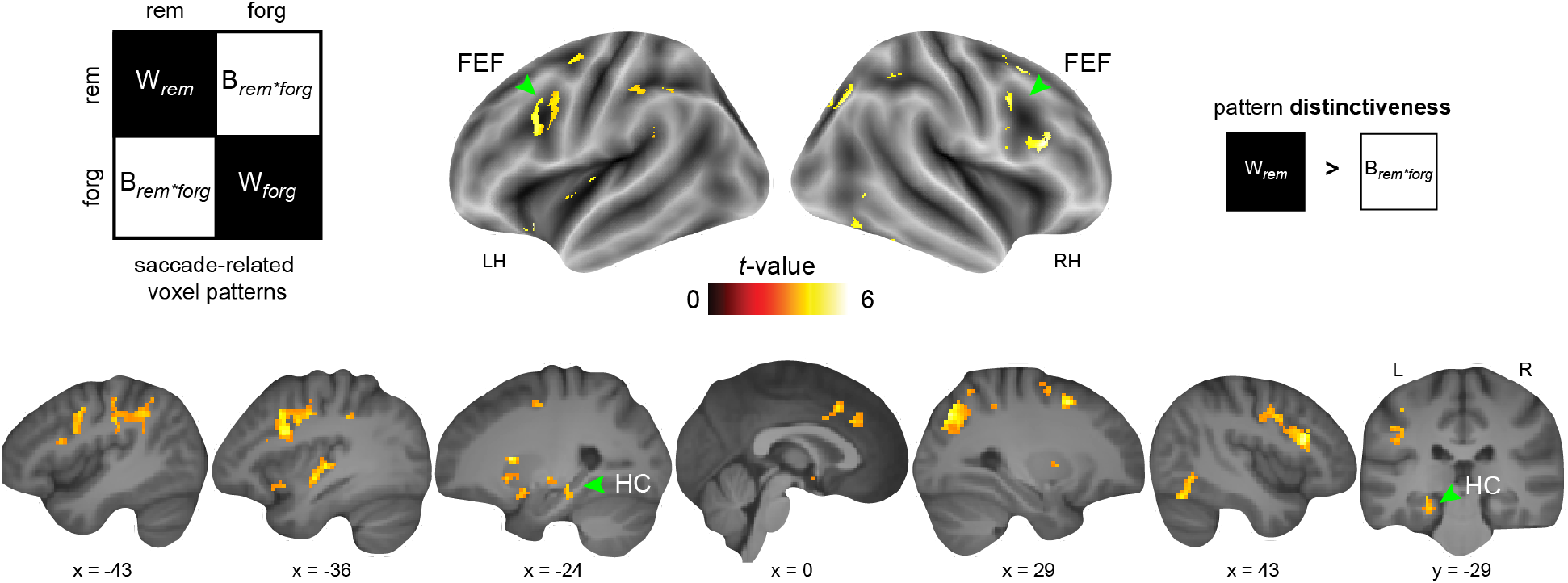
Saccade-related pattern distinctiveness predicts memory formation. The left upper panel shows a schematic of the correlation matrix for saccade-related representational similarity analysis (RSA). Searchlight-specific voxel patterns were sorted according to memory (rem, forg) and average pattern similarity was obtained for within-(W_*rem*_, W_*forg*_) and between-condition quadrants of the matrix (B_*rem*forg*_). Results show saccade-related pattern distinctiveness obtained through whole-brain, searchlight-based RSA. Results are shown at *p* < 0.05 FWE-corrected at cluster level (cluster-defining threshold of *p* < 0.001). LH, left hemisphere, RH, right hemisphere, L, left, R, right.

First, we found that saccade-related voxel patterns during the encoding of later remembered scenes did not appear significantly more stable compared to patterns during later forgotten scenes (i.e., no difference in within-condition pattern similarities; one-sample *t*-test, *N* = 32, contrast W_rem_ > W_forg_, not shown in figure). This was confirmed by additional ROI analysis, showing that saccades were associated with generally stable HC patterns, independent of whether material was later remembered or forgotten (**Results S2, Figure S5**).

Second, results showed that saccade-related patterns for later remembered scenes appeared distinct from patterns during later forgotten material in the left HC and the adjacent medial temporal lobe, the bilateral FEF, and the right superior parietal cortex (i.e., higher within-than between-condition pattern similarity; one-sample *t*-test, *N* = 32, contrast W_rem_ > B_rem*forg_; **Figure 4, Table S6**). This profile also included the dorsal anterior cingulate cortex and insula, the bilateral ventral striatum, as well as the right inferior temporal gyrus. Hence, while saccades were linked to stable patterns overall, saccade-related patterns during later remembered scenes appeared distinct from those during later forgotten scenes within in a wide-spread set of regions, including the HC and FEF.

### Saccade-related increase in hippocampal pattern distinctiveness is associated with stronger inhibition of the visual cortex during memory formation

Lastly, we reasoned that the two memory-related effects we found might be correlated: if the HC inhibits the VC during successful memory formation, this might be related to saccade-related HC patterns during later remembered scenes that are more distinct from those associated with later forgotten scenes (**Figure 5**). Testing this potential link, we performed correlation analysis between the strength of inhibitory HC-to-VC coupling (which was modulated by MEMORY, saccades_rem_ > saccades_forg_) and the average pattern distinctiveness across saccade-related volumes extracted from the left HC (contrast W_rem_ > B_rem*forg_, same left-lateralized ROI as for DCM analysis).

**Figure 5.**
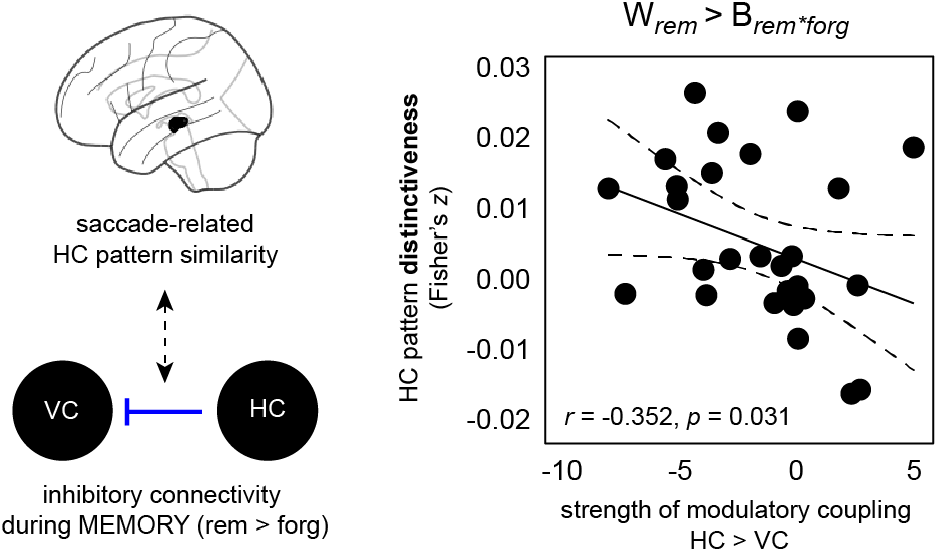
Saccade-related increase in hippocampal pattern distinctiveness is associated with stronger inhibition of the visual cortex during memory formation. Correlation between saccade-related inhibitory HC-to-VC coupling during successful memory formation (obtained from DCM analysis, left hemisphere) and saccade-related changes in hippocampal pattern similarity during successful memory formation (left hemisphere, same ROI as also for DCM analysis).

Across participants, stronger inhibition of the VC by the HC during successful memory formation was associated with more distinct saccade-related HC patterns during successful memory formation [*N* = 29, 3 formal outliers excluded, *r*_Pearson_ = -0.352, 95% CI = [-0.64, 0.02], *p*_*one-tailed*_ = 0.031; **Figure 5**]. In other words, as participants visually encoded scenes, HC patterns were more distinct for later remembered material and the strength of this effect positively scaled with the degree to which the HC inhibited the VC. This was not significant in the right hemisphere (*p*_*one-tailed*_ = 0.347), but note that the HC pattern similarity effects appeared mainly left-lateralized (**Figure 4**). Additional analyses confirmed that this relationship was specifically due to HC pattern distinctiveness rather than pattern stability (**Results S3, Figure S6**).

## Discussion

In this study, we asked which of the functional routes between the visual cortex, frontal eye fields, and the hippocampus facilitates memory during visual exploration and whether saccades affect neural representations. By combining network modeling and multivariate pattern analysis of saccade-related BOLD activity, we obtained several findings advancing the current understanding of how neuronal processes and saccades interrelate in the service of memory: The visual cortex (VC; and to a lesser extent the frontal eye fields, FEF) increased activity in the hippocampus (HC) as participants visually explored scenes, independent of memory. This general effect was complemented by saccade-related inhibition of VC activity by the HC that was stronger as participants viewed scenes that were later remembered (compared to later forgotten scenes). In terms of HC representations, we found that saccade-related HC patterns were stable overall (independent of subsequent memory), but that HC patterns during later remembered scenes appeared distinct from patterns during later forgotten material. Finally, the degree of HC pattern distinctiveness scaled with the strength of the saccade-related inhibitory HC-to-VC connection while viewing later remembered scenes. These findings highlight a functional coordination of neuronal signals and visual exploration behavior, and show saccade-related network dynamics and neural representations in a visuo-oculomotor-HC circuit that support memory formation.

We hypothesized that saccade-related directed connectivity between visuo-oculomotor (VC, FEF) and medial temporal regions (HC) supports memory. Using DCM, we found that the model where saccade-related activity drove the network via both the VC and FEF explained our data best (**Figure 2C**). Within this model, excitatory connectivity from the VC towards the HC (and to a weaker extent from the FEF towards the HC; **Figure 2D**) was most pronounced. Oculomotor control primarily involves the FEF but also the VC (Munoz and Everling, 2004), and the FEF is associated with the planning and generation of voluntary saccades (Connolly et al., 2005; Everling and Munoz, 2000; Johnston and Everling, 2008; Vernet et al., 2014; Wurtz et al., 2001). Visuo-oculomotor and HC regions are structurally connected through a distributed set of polysynaptic routes (Shen et al., 2016), setting the stage for a functional connection between the areas. In line with this notion, previous work demonstrated that saccades coordinate neuronal activity in the HC (Doucet et al., 2020; Hoffman et al., 2013; Jutras et al., 2013; Mao et al., 2021; Staudigl et al., 2022) and other medial temporal lobe regions (Ringo et al., 1994; Sobotka et al., 1997; Sobotka and Ringo, 1997), as well as the functional connectivity between regions (Sobotka et al., 2002; Staudigl et al., 2022). Our results support and expand these findings, showing that the VC (and to a lesser degree the FEF) drove HC activity by means of excitatory coupling when participants made saccades. We speculate that such enhanced coupling time-locked to eye movements might pave the way for inter-regional information transfer (Buzsaki and Draguhn, 2004; Fries, 2005; Leszczynski and Schroeder, 2019), perhaps re-setting the phase of low-frequency oscillations (Voloh and Womelsdorf, 2016) to facilitate HC processing (Liu et al., 2017; Staudigl et al., 2022) and the prediction of upcoming visual input (Henderson, 2017; Renninger et al., 2007). Moreover, we found that bilateral functional connectivity between the VC and FEF coincided with saccade production (**Figure 2D**). The FEF directly project towards the primary VC and extrastriate regions (Barone et al., 2000; Markov and Kennedy, 2013). FEF stimulation was shown to modulate neuronal firing in visual areas (Armstrong et al., 2006; Moore and Armstrong, 2003) presumably to guide visual attention (Michalareas et al., 2016; Popov et al., 2017; Veniero et al., 2021; Wang et al., 2016). Excitatory FEF-to-VC coupling could thus influence processing in early visual regions and, in return, might be regulated by inhibitory VC-to-FEF signals (**Figure 2E**). Altogether, we found that excitatory coupling from visuo-oculomotor regions towards the HC was time-locked to saccades but that it appeared independent of whether content was later remembered or forgotten. This suggests that a general saccade-related connectivity change at the time of stimulus encoding might be complemented by other, memory-specific processes.

In a next step, we tested which of the saccade-related functional routes within the VC-FEF-HC circuit might be modulated by memory formation. Results revealed saccade-related inhibitory coupling from the HC towards the VC, showing that HC activity decreased VC activity more strongly as participants visually explored scenes that were later remembered (**Figure 2F**). This partly fits with a recent report by Kragel and colleagues who investigated HC oscillations that were recorded invasively from human patients as these made eye movements to perform a spatial memory task. The authors discovered that phase-locking at the peak of HC theta oscillations preceded eye movements towards remembered locations, indicating that HC processing might modulate memory-guided viewing behavior (Kragel et al., 2020). Furthermore, HC theta was shown to exert top-down influence over visual attention networks as participants made eye movements to encode novel scenes (Kragel et al., 2021), and was shown to modulate V1 responses as rodents navigated a familiar spatial environment (Fournier et al., 2020). Considering these results, our finding of saccade-related HC-to-VC coupling during memory formation supports an interpretation in which (memory) signals might inhibit early visual processing and inform the location of upcoming saccades, in agreement with a theoretical account of predictive coding (Hindy et al., 2016; Kok and Turk-Browne, 2018; Wynn et al., 2020). For example, when encoding a novel scene, neural representations in the medial temporal lobe might provide a “mental framework” for the current experience by reconciling incoming perception with recent input (e.g., based on the recent saccade history) and prior knowledge (e.g., based on previous experiences with similar scenes) within repeated encoding-retrieval cycles (see also Kragel and Voss, 2022). Such recurrent processing would enable predictions concerning the location of relevant features within the scene and could hence guide future eye movements. This might involve visually responsive cells in the medial temporal lobe that were shown to code for saccade direction (Killian et al., 2015), gaze position (Meister and Buffalo, 2018), and that provide a map of visual space in non-human (Killian et al., 2012) and human primates (Nau et al., 2018; Staudigl et al., 2018), potentially aiding visual recognition memory (Bicanski and Burgess, 2019). Taken together, saccade-related connectivity changes within the VC-FEF-HC network are in part modulated by memory, showing HC inhibition of the VC as participants visually explored later remembered scenes.

Saccades move the fovea across a visual scene to enable detailed visual analysis of the particular location that is subsequently fixated. Put differently, the function of saccades could be to direct overt attention towards to-be-encoded features of a visual scene and thereby influence which information will be committed to memory. One possibility to characterize the associated neural dynamics is by focusing on the activity within (local) voxel patterns and to quantify their similarity across cognitive processes. For example, memory formation was previously linked to pattern similarity in medial temporal and neocortical regions (Aly and Turk-Browne, 2016; LaRocque et al., 2013; Wagner et al., 2016; Xue et al., 2013). Aly and Turk-Browne investigated the effect of attention on HC patterns by manipulating participants’ attentional state, asking them to switch focus between different aspects while keeping visual input constant. HC patterns showed greater similarity (or “stability”) across trials of the same attentional state compared to trials of a different attentional state (Aly and Turk-Browne, 2015). The authors then went on to show that HC pattern stability at encoding promoted memory and that this was paralleled by multivariate connectivity with neocortical regions in which pattern stability was also affected by attention (Aly and Turk-Browne, 2016). What remained unclear, however, was whether the effect of increased pattern stability could actually be tied to eye movements which are thought to reflect allocation of overt attention. To our surprise, we could not confirm previous results but instead discovered that saccade-related HC patterns were generally stable, independent of memory formation. However, when participants produced saccades to study scenes that were later remembered, patterns in the HC (and in a wide-spread set of regions, including the FEF) appeared distinct from saccade-related patterns when participants studied scenes that were later forgotten (**Figure 4**). This suggests that the saccade-related pattern stability is generally high and complemented by additional processes that disambiguate remembered from forgotten material (LaRocque et al., 2013; Liu et al., 2022). Moreover, pattern “distinctiveness” went hand-in-hand with stronger memory-modulated HC-to-VC inhibition across participants (**Figure 5**). We speculate that a transient saccade-related suppression of VC activity by the HC might foster memory formation through the integration of incoming sensory input (e.g., via the VC-to-HC and FEF-to-HC routes) with previously foveated content, thereby stitching together a stable percept of the respective scene image. Alternatively, suppressed activity in the VC could reflect more efficient visual processing by accounting for expectations about the visual scene that were generated by the HC (Kok et al., 2012; Lee and Mumford, 2003). Such sharpened activity in VC could be reflected in more distinct HC patterns during successful memory formation, in line with the correlational relationship we describe here. Speculatively, pattern similarity effects could be tied to the phase-reset of HC oscillations as was previously reported in monkeys (Jutras et al., 2013). By realigning the oscillatory phase, saccades might guide the organization of functional networks (Voloh and Womelsdorf, 2016) and form coherent but distinct neural patterns that promote memory formation and successful recognition (Aly and Turk-Browne, 2016, 2015; Brunec et al., 2020; LaRocque et al., 2013; Liu et al., 2022; Wagner et al., 2016). We encourage future research to examine a potential relationship between oscillatory phase-reset and HC pattern-based representations.

Besides the HC, we found that saccade-related pattern distinctiveness was predictive of subsequent memory in the FEF, the adjacent lateral prefrontal cortex and parietal regions (**Figure 4**). This set of areas resembles the so-called frontoparietal network which is thought to coordinate attention (Szczepanski et al., 2013; Vincent et al., 2008) and to promote memory formation (Cabeza et al., 2008; Ciaramelli et al., 2008; van Buuren et al., 2019). Furthermore, increased frontoparietal activity was associated with stronger pattern similarity in other regions predictive of subsequent memory (Xue et al., 2013) and pattern similarity was enhanced through stimulation of the lateral prefrontal cortex (Lu et al., 2015). Regarding the VC-FEF-HC circuit, we postulate that the FEF controls voluntary saccades during visual scene exploration (Connolly et al., 2005; Everling and Munoz, 2000; Johnston and Everling, 2008), possibly affecting pattern similarity and routing information to visually-responsive neurons in the medial temporal lobe that map visual space (Killian et al., 2012). The precise connectivity profile likely depends on the task at hand. For instance, saccades were shown to modulate neuronal firing in the parahippocampal gyrus during visual processing (Andrillon et al., 2015; Ringo et al., 1994; Sobotka et al., 1997), in the anterior thalamus during natural scene viewing (Leszczynski et al., 2021), in the amygdala as participants viewed faces (Minxha et al., 2017; and see also Staudigl et al., 2022), and in the superior temporal sulcus as participants viewed objects (Bartlett et al., 2011). Although we think that the VC-FEF-HC circuit is suitable for capturing the general processes associated with saccade-related memory formation, we acknowledge that our work only provides a first step and hope that future studies expand our findings by using larger-scale models.

On a final note, we did not find evidence for saccade-related HC-to-FEF connectivity. This contrasts with a recent report and modeling work by Ryan and colleagues on information flow between oculomotor and medial temporal regions, showing that HC stimulation evoked responses in the FEF (Ryan et al., 2020) and that making eye movements during a mental construction task (as compared to restricting them) led to increased HC-to-FEF connectivity (Ladyka-Wojcik et al., 2022). Differences in the paradigms and analysis strategies used could explain the divergence. By combining a fast MRI acquisition sequence with saccade-related fMRI analyses, we investigated the coordination of saccades and brain activity related to successful memory formation in a subsequent memory paradigm. Ladyka-Wojcik and colleagues (2022), on the other hand, compared free vs. restricted viewing conditions during a mental reconstruction task. On a more general level, the circuit dynamics might also change depending on the inclusion or exclusion of specific seeds and their exact placement, raising the possibility that saccade-related HC-to-FEF connectivity could differ for specific HC subfields or adjacent regions in the medial temporal lobe (as was also shown by Ryan et al., 2020).

In conclusion, we found that eye movements affected the direction of information flow within a visuo-oculomotor and medial temporal network. These saccade-related changes in functional connectivity at the time of encoding were complemented by other, memory-specific processes that determined whether participants would later remember or forget. Most importantly, saccades were linked to stable but distinct HC representations that were accompanied by stronger HC inhibition of the visual system during memory formation. We suggest a coordination of saccades and directed circuit dynamics via the visuo-oculomotor system to improve HC processing, thus setting the stage for memory formation.

## Acknowledgements

The study was performed at the Donders Institute for Brain, Cognition and Behaviour. The authors would like to thank Paul Gaalman for advice and assistance with MRI scanning and the eye-tracker setup.

## Author contributions

TS, CFD and OJ conceptualized the study; TS collected the data; ICW analyzed the data and wrote the first version of the manuscript; ICW and TS edited the manuscript; TS supervised the study; all authors discussed and interpreted the results and commented on the manuscript.

## Materials and Methods

### Participants

Forty-eight participants initially volunteered for this study. Of those, 16 participants were excluded due to not completing the study (7 individuals), excessive motion artefacts (4 individuals), technical problems during the data recording (3 individuals), a low number of identified saccades during the fMRI session (1 individual, < 30 detected saccades per condition), and a low memory recognition performance during the MRI session (1 individual, higher rate of false alarms than correct rejections). The final sample thus consisted of 32 participants (23 females, age range 18-30 years, mean age = 23 years, 32 right-handed). All individuals were healthy and did not report any history of neurological and/or psychiatric disorders, had normal or corrected-to-normal vision, and provided written informed consent before the start of the experiment. The study was reviewed and approved by local ethics committee (Commissie Mensgebonden Onderzoek, region Arnhem-Nijmegen, The Netherlands; CMO-2014/288).

### Study setup and task design

This study was part of a larger project investigating the effects of saccadic eye movements on memory processing using magnetencephalography (MEG) and magnetic resonance imaging (MRI). In brief, participants were invited to the laboratory on separate days and underwent either MEG (not reported here, but see Staudigl et al., 2017) or MRI while completing a recognition memory task. The session order was balanced across participants and the task comprised different stimulus sets for each session.

### Recognition memory task

During the study period, participants were instructed to memorize 200 photographs of unique indoor and outdoor scenes (100 each). Photographs were resized to a dimension of 1024 × 768 pixels and were presented on a black background. Each scene was shown for 4 s during which participants could freely view the photograph (thus, participants were not instructed to fixate). To ensure attention to each scene, participants were asked to judge whether the photograph depicted an indoor or outdoor scenario via button press during the subsequent fixation period (2125, 4125, or 7125 ms, 80/80/40 distribution across the 200 trials, pseudo-randomized), after which the next scene appeared. The order of scenes was pseudorandomized with no more than four scenes of the same type (indoor/outdoor) shown consecutively. The encoding period was followed by a distractor task (i.e., solving simple mathematical problems, 1 min) and a rest period (3 min).

During the recognition memory test period, participants viewed all scenes that were shown during the previous study period, intermixed with 100 novel scenes (half of them indoor/outdoor; i.e., a total of 300 scene photographs were presented). The assignment of scene photographs to study or recognition memory test periods was counterbalanced across participants. During the recognition memory test period, scenes were presented for 4 s each, followed by a 6-point rating scale that required participants to indicate whether they recognized the scene as “old” or “new” [self-paced; the scale ranged from “very sure old” (1) to “very sure new” (6)], and a fixation period until the next trial started (same as above; 2125, 4125, or 7125 ms, 80/80/40 distribution across the 200 trials, pseudo-randomized). The recognition memory test was divided into 2 blocks, with a short break in between. Participants were allowed to rest between blocks, if needed. After the recognition memory test, participants were asked to fixate on different locations on the screen to evaluate eye tracker accuracy (5 min), followed by the structural scan.

### Recognition performance (*d*-prime)

Based on individual performance during the recognition memory test period, trials were grouped into four bins: (1) scenes that were correctly judged at “old” (i.e., hits, collapsing across ratings 1-3, mean ± standard error of the mean (s.e.m.): 140.9 ± 5.6 trials); (2) scenes that were correctly judged as “new (i.e., correct rejections, collapsing across ratings 4-6, 85.8 ± 1.8 trials); (3) scenes that were incorrectly judged as “old” (i.e., false alarms, collapsing across ratings 1-3, 14.2 ± 1.8 trials); (4) scenes that were incorrectly judged as “new” (i.e., misses, collapsing across ratings 4-6, 59.1 ± 5.6 trials). None of the participants displayed any actually missed trials without button presses. Individual hit and false alarm rates were *z*-scored and recognition performance (*d*-prime) was calculated as [*z*(hits) – *z*(false alarms)].

### Eye tracking data acquisition, analysis, and saccade detection

To capture saccadic eve movements, we recorded horizontal and vertical eye gaze, as well as pupil size using a video-based infrared eye tracker (EyeLink 1000 Plus, SR Research, Ontario, Canada). Before recording, raw eye movement data was mapped onto screen coordinates by means of a calibration procedure. Participants sequentially fixated on nine fixation points on the screen, arranged in a 3 × 3 grid. This was followed by a validation procedure during which the nine fixation points were presented once more while the differences between the current and previously obtained gaze fixations (from the calibration period) were measured. If these differences were < 1° of visual angle, the calibration settings were accepted and the eye tracker recording was started.

Eye tracking data was processed using Fieldtrip (https://www.fieldtriptoolbox.org). Saccadic eye movements were identified by transforming vertical and horizontal eye movements into velocities, whereby velocities exceeding a threshold of 6 × the standard deviation (s.d.) of the velocity distribution and with a duration of > 12 ms were defined as saccades (Engbert and Kliegl, 2003). Saccade onsets during trials of the study period (i.e., during the presentation of scene photographs) were defined as events-of-interest. Only saccades that followed a minimum fixation period of 25 ms were included. Saccades that were followed or preceded by blinks (+/-100 ms) were excluded (blinks were defined as large deflections in pupil diameter: mean ± 5 standard deviations; eye tracking data in the vicinity of blinks is unreliable due saturation effects). Trials with more than 25% of missing eye tracker data were discarded. We detected a total of 48510 saccades in the eye tracking data (*N* = 32; average number of saccades per participant, mean ± SEM: 1515.94 ± 80.95 saccades).

### MRI data acquisition

Imaging data were collected at the Donders Institute for Brain, Cognition and Behaviour (Nijmegen, The Netherlands), using a 3T Prisma Fit scanner (Siemens, Erlangen, Germany) equipped with a 32-channel head coil. We acquired on average 2456 (± 5.3) T2*-weighted blood oxygen level-dependent (BOLD) images during the study period of the recognition memory task, using the following echo-planar imaging (EPI) sequence: repetition time (TR) = 657 ms, echo time (TE) = 30.8 ms, multi-band acceleration factor = 8, 72 axial slices, interleaved acquisition, field of view (FoV) = 174 × 174 mm, 72 × 72 matrix, flip angle = 53°, slice thickness = 2.4 mm, no slice gap, voxel size = 2.4 mm isotropic. The structural image was acquired using a standard magnetization-prepared rapid gradient-echo (MPRAGE) sequence with the following parameters: TR = 2300 ms, TR = 3.03 ms, FoV = 256 × 256 mm, flip angle = 8°, voxel size = 1 mm isotropic.

### MRI data preprocessing

The fMRI data were processed with SPM8, GLM model comparison and DCM analysis were performed with SPM12 (http://www.fil.ion.ucl.ac.uk/spm/) in combination with MATLAB (The Mathworks, Natick, MA, USA). The first 12 volumes were excluded to allow for T1-equilibration. The remaining volumes (of both the encoding and recognition periods) were realigned to the mean image. The structural scan was co-registered to the mean functional image and was segmented into grey matter, white matter, and cerebrospinal fluid using the “New Segmentation” algorithm. All images (functional and structural) were then spatially normalized to the Montreal Neurological Institute (MNI) EPI template using Diffeomorphic Anatomical Registration Through Exponentiated Lie Algebra (DARTEL; Ashburner, 2007), and functional images were further smoothed with a 3D Gaussian kernel (6 mm full-width at half-maximum, FWHM).

### fMRI data modeling

We investigated subsequent memory effects by sorting all scenes presented during the study period (or all saccades) based on individual memory performance during the recognition test period. In other words, scenes (or saccades during scenes) that were correctly judged as “old” (i.e., hits) or “new” (i.e., correct rejections) during the recognition test period were defined as “remembered” (*N* = 32; average numbers per participant, mean ± SEM: 226.7 ± 5.4 scenes, 1140.25 ± 85.34 saccades). Scenes (or saccades during scenes) incorrectly judged as “old” (i.e., false alarms) or “new” (i.e., misses) were defined as “forgotten” (73.3 ± 5.4 scenes, 375.69 ± 39.13 saccades).

### Saccade-related model

To be able to capture memory-related activation changes associated with saccadic eye movements, we employed a fast MRI acquisition sequence (TR = 657 ms) that allowed us to identify brain volumes during which at least one saccade appeared (*N* = 32; average numbers of volumes per participant, mean ± SEM; total: 861.8 ± 38.4 volumes; remembered: 633 ± 40.2 volumes; forgotten: 228.8 ± 25.8 volumes; average number of saccades per volume; total: 1.75 ± 0.03 saccades; remembered: 1.77 ± 0.03 saccades; later forgotten: 1.69 ± 0.03 saccades). On average, each trial (i.e., scene) was associated with 7.3 (± 0.4) volumes with at least one saccade. Related to the higher average number of saccades during remembered trials (see above), there was also a higher average number of saccade-related brain volumes during remembered (7.95 ± 0.4 volumes) compared to forgotten trials (6.65 ± 0.43 volumes; *t*(31) = 8.26, Cohen’s *d* = 1.46, 95% CI = [0.99, 1.63], *p*_two-tailed_ < 0.001).

These brain volumes defined our events-of-interest, were incorporated into the General Linear Model (GLM) framework, and were separated into two task regressors that described the effects of saccades during later remembered and later forgotten trials (saccades_rem_, saccades_forg_). The BOLD responses of these regressors were modelled with a stick function (0 s duration). General task effects were modelled with a task regressor time-locked to the presentation onset of each scene photograph and were estimated as a boxcar function (4 s duration). All events were convolved with a standard canonical hemodynamic response function (HRF). Additionally, the six realignment parameters, their first derivatives, and the squared first derivatives were included in the design matrix. This resulted in 18 additional regressors that accounted for noise due to head movement. A high-pass filter with a cutoff at 128 s was applied and individual contrast maps were created ([saccades_rem_ ∩ saccades_forg_]> baseline, saccades_rem_/saccades_forg_ > baseline, saccades_rem_ vs. saccades_forg_). Group effects were investigated by submitting contrast images to separate one-sample *t*-tests.

### Parametric and conventional models

We estimated two additional GLMs to determine whether the abovementioned saccade-related model provided a better fit to the fMRI data during the study period of the recognition memory task as compared to other, more conventional models. These models differed only in the arrangement of the task-specific regressors; regressors capturing head motion, filter specifications, and the statistical testing for group effects were identical to above.

First, using a parametric approach, we modelled the BOLD response for all trials with separate task regressors time-locked to the onset of each scene photograph (scene_rem_, scene_forg_). These events were estimated as a boxcar function with the duration of one trial (4 s) and were convolved with a canonical HRF. Parametric modulators for each of these regressors captured the number of saccades per trial. Contrast maps were created to detect the effect of memory (scene_rem_ > scene_forg_) as well as effects parametrically modulated by the number of saccades (scene_rem_saccades_ > scene_forg_saccades_).

Second, to compare effects to previous studies that tested for subsequent memory effects but did not take into account eye movements, we defined a conventional model that was oblivious to saccadic events. Again, the BOLD response for all trials was modelled with separate task regressors time-locked to the onset of each scene photograph (scene_rem_, scene_forg_), estimated as a boxcar function with the duration of one trial (4 s), convolved with a canonical HRF, and tested using the contrast scene_rem_ > scene_forg_.

### Model comparison

To verify that the saccade-related model provided a better model fit to our fMRI data during the study period than the parametric model we performed model comparison using the SPM toolbox for Model Assessment, Comparison and Selection (MACS; Soch and Allefeld, 2018). We did not include the conventional model in this comparison since it did not take into account saccade-related activation changes.

Following the procedures described previously (Soch and Allefeld, 2018), we first defined a model space that comprised the saccade-related and parametric GLMs of each participant. Model assessment (i.e., the quantification of relative model quality) was performed by calculating the Bayesian cross-validated log model evidence (LME). This measure takes into account model accuracy as well as complexity and circumvents the problem of defining prior distributions on the model parameters by obtaining priors based on a part of the available data, calculating the LME on the remaining data (Soch et al., 2016). This resulted in participant- and model-specific LME maps that quantified the voxel-wise model performance, followed by model comparison (i.e., the quantification of the difference in model quality between models) using the log Bayes factor (LBF) and the posterior model probabilities (PP). Model selection was performed to determine the model that best explained the data (i.e., the model with the highest estimated model frequency, quantified by the likeliest frequency, LF) by employing group-level, random-effects Bayesian model selection (BMS). This approach accounts for the fact that different models might be optimal in different participants and shows how frequently each model is optimal in the given participant population. BMS resulted in participant-specific whole-brain maps that captured the voxel-wise estimated model frequencies (LF) and exceedance probabilities.

Finally, the BMS outcomes were used to calculate selected-model maps (SMMs, one for each model) that indicated the model with the highest frequency per voxel. Using these SMMs, we (1) calculated the percentage of voxels across the whole brain that favored a respective model, and (2) directly assessed model fit in the three regions-of-interest (see below; i.e., we extracted the average, model-specific LF values for the left and right visual cortex, hippocampus, and the frontal eye fields).

### Definition of regions-of-interest (ROIs)

Regions-of-interest (ROIs) for the visual cortex (VC), the frontal eye fields (FEF), and the hippocampus (HC) were defined based on results from the group statistical analysis (saccade-related model). The visual cortex was defined using the contrast [saccades_rem_ ∩ saccades_forg_]> baseline (thus, capturing the general effect of saccadic eye movements irrespective of subsequent memory; **Figure 1D**). Results were masked with a bilateral anatomical image of the visual cortex provided by the Automatic Anatomical Labeling (AAL) atlas (Tzourio-Mazoyer et al., 2002) and the location of the peak activation value within this selection was determined (left VC: *x* = -7, *y* = -94, *z* = -7). We then placed a sphere (15 mm diameter) around the coordinate and once again masked it with the abovementioned bilateral anatomical image to assure that only voxels within the anatomical borders of the visual cortex were included in the ROI.

The FEF and HC were defined using the contrast saccades_rem_ > saccades_forg_ (**Figure 1E**). Results were again masked with a bilateral anatomical image of the HC obtained from the AAL atlas. The coordinate associated with the peak activation value was determined (right HC: *x* = 36, *y* = -29, *z* = -12), was surrounded with a 15 mm sphere, and was masked once more with the bilateral anatomical image. In accordance with previous work investigating oculomotor control (Johnston and Everling, 2008; Munoz and Everling, 2004), the FEF was defined through the cluster peak coordinate (right FEF: *x* = 43, *y* = 7, *z* = 29) and was surrounded with a 10 mm sphere. The three ROIs were mirrored to create a contralateral counter-part for each ROI and voxels that overlapped between the left and right visual cortex were excluded from both masks (left/right VC = 151 voxels; left/right HC = 71 voxels; left/right FEF = 160 voxels).

## Connectivity analyses

### Dynamic Causal Modeling (DCM)

We used dynamic causal modeling (DCM; Stephan and Friston, 2010) to gauge the effective connectivity (i.e., the directional interactions) within a ROI-based network that included the visual cortex, the hippocampus, and the FEF. We initially focused on the left-lateralized network and subsequently validated the results using the right-lateralized network, probing that both analyses yielded virtually identical outcomes (see **Figure 3, Table S4**).

First, we adapted the saccade-related model (see above) to test directional network interactions that were driven by saccadic eye movements and modulated by subsequent memory (yielding a separate GLM_DCM_). We specified a regressor marking brain volumes during which one or more saccades occurred (SACCADE, 0 s duration), a parametric modulator that captured whether a given saccade appeared during a trial that was later remembered or forgotten (MEMORY, saccades_rem_ > saccades_forg_), as well as a regressors that captured the general effect of task (i.e., scene presentations; TASK, 4 s duration). We then extracted the first eigenvariate of each ROI’s functional timecourse and adjusted it for average activation levels using an *F*-contrast (visual cortex: timecourse associated with SACCADE, hippocampus and FEF: timecourse associated with MEMORY). To ensure that we selected voxels linked to task-related activity fluctuations rather than noise, timecourse extraction was based on a threshold of *p* < 0.05 (uncorrected), and ROI images were additionally masked with the participant-specific brainmask to exclude out-of-brain voxels (which was based on the overlap of each participant’s EPI-based, normalized field-of-view with the standard-space brainmask provided by SPM).

The DCM model space was created as follows (see also Zeidman et al., 2019): (1) all models were fully connected (i.e., bilateral connections between all 3 ROIs) with self-connections (matrix *A*); (2) modulatory effects (i.e., later remembered/forgotten, MEMORY) could act on the intrinsic (i.e., between-region) connections (matrix *B*); and (3) driving inputs were given through SACCADE that could drive the visual cortex, FEF, or both (matrix *C*). Overall, the model space thus included 192 models that were grouped into 3 families, depending on the location of the SACCADE driving input. We did not modulated the self-connections as this would have substantially enlarged our model space.

All models were estimated for each participant and group-level DCM inference was performed in two steps. First, we performed Bayesian Model Selection to reduce the model space and to define the winning model family (Penny et al., 2004). This described a group of models for which the regressor SACCADE was driving model effects with the highest model evidence via (a) specific region(s) (i.e., via the visual cortex, FEF, or both), irrespective of further assumptions about the model structure. Second, we performed random-effects Bayesian Model Averaging (BMA) within the winning model family to obtain average parameter estimates that were weighted by the respective posterior probability of the models in the model space (Penny et al., 2010). This resulted in a final group model that took uncertainty about the specific model structure into account (Stephan et al., 2010). One-sample *t*-tests were performed to test for significance of intrinsic or modulatory MEMORY connections (applying Bonferroni correction for multiple comparisons).

### Psychophysiological Interaction (PPI) analysis

Next, we performed psychophysiological interaction analysis (PPI; Friston et al., 1997) to uncover the brain-wide connectivity profiles of the VC, FEF and HC during saccade-related memory encoding rather than just focusing on their local connectivity within the confined DCM network. To achieve this, we took the seed regions obtained from our initial analysis (left VC, right FEF and right HC, see above) and extracted the first eigenvariate of each regions’ functional timecourse, adjusted for average activation levels using an *F*-contrast (taking the original, saccade-related model). The timecourse was then deconvolved to estimate the putative neural activity of the seed region (i.e., the physiological factor) and was multiplied with stick functions that defined the specific task events (i.e., the psychological factor). The resulting vectors were convolved with the canonical HRF yielding one PPI regressor per condition-of-interest, and contrasts were created (saccades_rem_ > saccades_forg_). Group-level connectivity analyses were performed using a set of one-sample *t*-tests.

### Representational Similarity Analysis (RSA)

Finally, we tested whether saccades affected neural representations depending on subsequent memory performance by focusing on changes in pattern similarity within a given voxel pattern. We reasoned that if saccades affected hippocampal processing to allow for successful memory formation, later remembered (compared to forgotten) trials should be linked to increased pattern similarity in brain regions associated with visual processing, oculomotor control (Johnston and Everling, 2008; Munoz and Everling, 2004) and memory formation (Aly and Turk-Browne, 2016, 2015; Kim, 2011; Spaniol et al., 2009), indexing pattern “stability”. Additionally, voxel patterns during memory formation might be distinct from those during the encoding of later forgotten material (indexing pattern “distinctiveness”), which might aid subsequent recognition (LaRocque et al., 2013).

We moved a spherical searchlight with a radius of 8 mm (171 voxels) throughout the brain volume (Kriegeskorte et al., 2006; Wagner et al., 2020, 2016), considering only searchlights that contained at least 30 gray matter voxels. Saccade-specific beta images (i.e., beta images for single brain volumes/TRs that contained at least one saccade) were obtained by defining and estimating a separate, saccade-specific GLM (GLM_RSA_) that contained as many saccade-related regressors as there were brain volumes with at least one saccade (Mumford et al., 2014), as well as regressors to capture general task effects and variations in head motion (identical to the original, saccade-related model). Single-volume, saccade-specific beta images were then contrasted against the implicit baseline and were averaged within a given trial.

Using the resulting *t*-maps, voxel-specific values were extracted for a given searchlight, data was reshaped into a trial × voxel matrix, and trials were sorted according to the two conditions (i.e., saccades_rem_, saccades_forg_). Data were *z*-scored across saccades to remove mean activation differences, and voxel patterns of each trial were correlated with the voxel patterns of all other trials, resulting in a trial × trial similarity matrix. This matrix was then Fisher’s *z*-transformed and pattern similarity scores were calculated by averaging across the respective quadrants of the similarity matrix. We then extracted the average pattern similarity separately across all later remembered and forgotten trials (within-condition similarity, W_rem_ and W_forg_, respectively) as well as the average pattern similarity across all later remembered and forgotten trials (between-condition similarity, B_rem*forg_). The resulting pattern similarity values were then assigned to each searchlight’s center voxel, yielding three 3-dimensional whole-brain pattern similarity maps per participant. Group-level significance was tested with a paired-sample *t*-test, comparing within-to within-(contrast W_rem_ > W_forg_), as well as within-to between-condition pattern similarity (contrast W_rem_ > B_rem*forg_).

### Statistical analysis of behavioral data and ROI-based fMRI results

Analyses of recognition performance (*d*-prime) and ROI-based fMRI results (coupling strength of DCM parameters, pattern similarity) were carried out with R (https://www.r-project.org) using a set of *t*-tests and correlations. Effect sizes were calculated as Cohen’s *d* (Lakens, 2013). As mentioned above, we *a priori* expected that modulatory coupling strength would be associated with increases in both hippocampal pattern stability and distinctiveness, which is why we adopted an α-level of 0.05 (one-tailed). For DCM analysis, we applied Bonferroni-correction to account for multiple comparisons when comparing driving inputs (α_Bonferroni_ = 0.05/2 driving inputs = 0.025), intrinsic connections (α_Bonferroni_ = 0.05/9 connections = 0.006), and modulatory connections (α_Bonferroni_ = 0.05/6 connections = 0.008). For all other cases, the α-level was set to 0.05 (two-tailed). Any exploratory analyses are explicitly described as such. Values from ROI-based fMRI analysis (coupling strength of DCM parameters, pattern similarity) that exceeded the median value ± 3 × the median absolute deviation were excluded from the analyses.

### Statistical thresholding of whole-brain fMRI results and anatomical labeling

Unless stated otherwise, significance for all whole-brain fMRI analyses was assessed using cluster-inference with a cluster-defining threshold of *p* < 0.001 and a cluster-probability of *p* < 0.05 family-wise error (FWE) corrected for multiple comparisons. The corrected cluster size (i.e., the spatial extent of a cluster that is required in order to be labeled as significant) was calculated using the SPM extension “CorrClusTh.m” and the Newton-Raphson search method (script provided by Thomas Nichols, University of Warwick, United Kingdom, and Marko Wilke, University of Tübingen, Germany; http://www2.warwick.ac.uk/fac/sci/statistics/staff/academic-research/nichols/scripts/spm/).

Anatomical nomenclature for all tables was obtained from the Laboratory for Neuro Imaging (LONI) Brain Atlas (LBPA40, http://www.loni.usc.edu/atlases/).

## Data and code availability

All anonymized data are available upon reasonable request to the authors in accordance with the requirements of the institute, the funding body, and the institutional ethics board. All analyses are based on openly available software. Raw data (recognition memory test performance, saccade-related data), DCM and RSA results, as well as (un-)thresholded statistical whole-brain fMRI maps and their corresponding results tables are available at the Open Science Framework (https://osf.io/4dshu/).

## Supplementary Materials

### Results S1: Model comparison in visual cortex, frontal eye fields, and hippocampus

To obtain a measure of how well the saccade-related model fitted voxel responses in the visual cortex (VC), frontal eye fields (FEF), and the hippocampus (HC), we extracted the average likeliest frequency (LF) from the respective ROIs using the selected-model maps (SMMs) that we had obtained from Bayesian Model Selection (**Figure S3, Materials and Methods**). The saccade-related model provided better model fit in almost all of the left- and right-lateralized ROIs (average LF for saccade-related model: left VC, 0.87; right VC, 0.73; right FEF, 0.64; left HC, 0.88; right HC: 0.79; no average LF values for the parametric model as it did not fit voxel responses in these regions). For the right FEF, both models provided comparable model fit (average LF: saccade-related model, 0.57; parametric model: 0.54).

### Results S2: Saccade-related HC pattern stability

In our whole-brain analysis, we found that saccade-related voxel patterns during the encoding of later remembered scenes did not appear significantly more stable compared to patterns during later forgotten scenes. We confirmed this with additional ROI-analysis: we focused on the HC (same left-lateralized ROI as for DCM analysis) and extracted the average within-condition pattern similarity values for later remembered and forgotten trials. Saccade-related patterns were generally stable (pattern similarity was significantly increased above zero, **Figure S5**; separate one-sample *t*-tests: W_rem_, *N* = 27, 5 formal outliers excluded, *t*(26) = 5.58, 95% CI = [0.01, 0.012], *p*_two-tailed_ < 0.0001; W_forg_, *N* = 21, 8 missing values, 3 formal outliers excluded, *t*(20) = 8.56, 95% CI = [0.01, 0.02], *p*_two-tailed_ < 0.0001) but there was no significant difference between the conditions (*p*_two-tailed_ = 0.077). Thus, saccades were associated with stable HC patterns, independent of whether material was later remembered or forgotten.

### Results S3: HC-to-VC inhibition is specifically related to distinct HC patterns during memory formation

Above, we had found that as participants visually encoded scenes, HC patterns for later remembered material were distinct from forgotten material and that the strength of this effect positively scaled with the degree to which the HC inhibited the VC. Additional analysis confirmed that this relationship was specifically due to HC pattern distinctiveness, showing (a trend towards) inhibitory HC-to-VC coupling that was lower when patterns were less distinct during memory formation (i.e., when the between-condition pattern similarity was higher, B_rem*forg_; *N* = 28, 4 formal outliers excluded, *r*_Pearson_ = 0.36, 95% CI = [-0.02, 0.54], *p*_two-tailed_ = 0.06; **Figure S6A**). There was no significant relationship between inhibitory HC-to-VC coupling and pattern stability as participants studied later remembered (W_rem_) or later forgotten (W_forg_) material (all *p*_two-tailed_ > 0.5, **Figure S6B**). Finally, there was no significant correlation between saccade-related excitatory VC-to-HC coupling and HC pattern distinctiveness or stability (all *p*_two-tailed_ > 0.05, not shown in figure).

## Supplementary Figures

**Figure S1:**
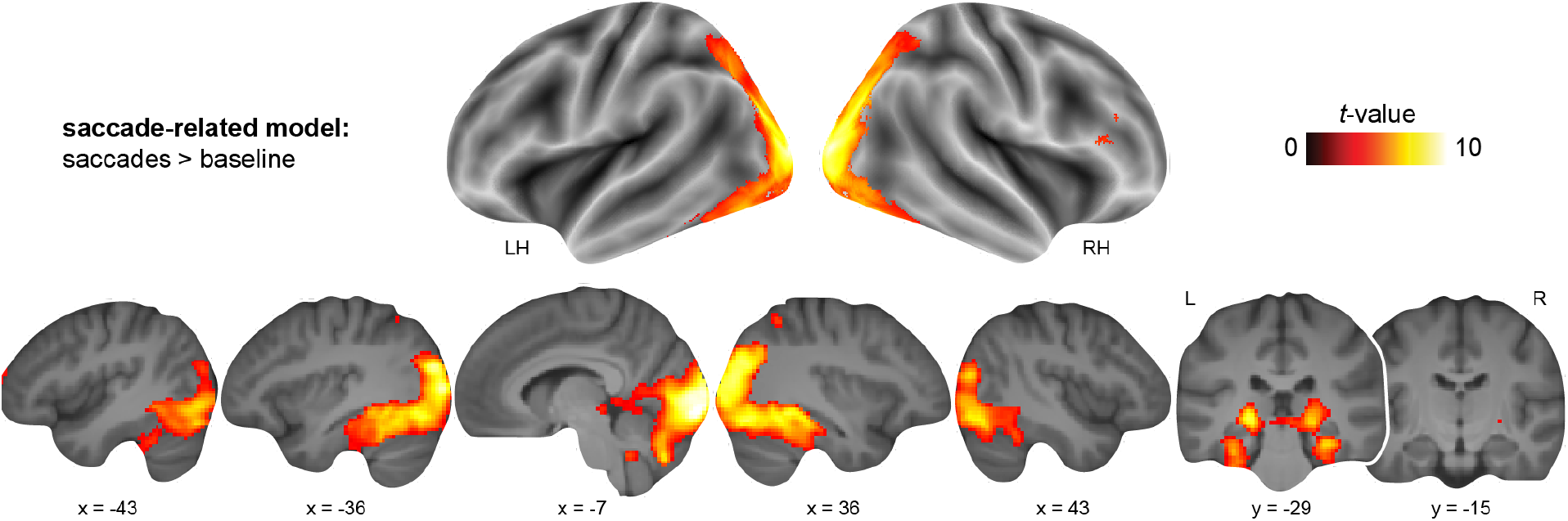
Saccade-related brain activation. Increased brain activation during saccadic eye movements (contrast [saccades_rem_ ∩ saccades_forg_] > baseline; **Table S1**), plotted onto brain slices of the average structural image. All results are shown at *p* < 0.05 FWE-corrected at cluster level (cluster-defining threshold of *p* < 0.001).

**Figure S2:**
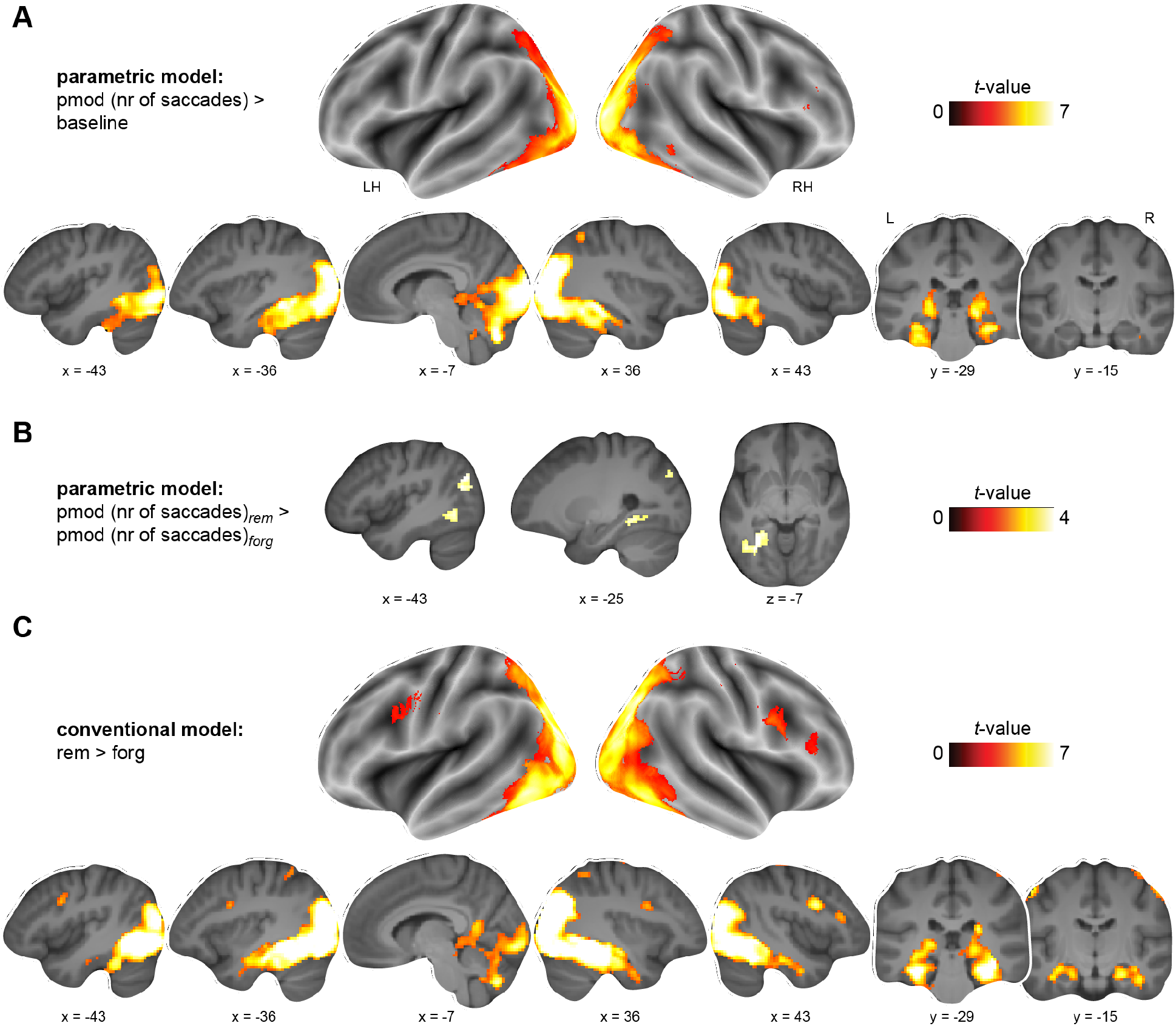
Brain activation using the parametric and conventional models. Parametric model: increased brain activation **(A)** at larger number of saccades during scene presentation (parametric modulator, pmod; contrast pmod (nr of saccades) > baseline; **Table S2**), and **(B)** at larger number of saccades during later remembered compared to forgotten scenes (contrast pmod (nr of saccades)_rem_ > pmod (nr of saccades)_forg_; **Table S2**). **(C)** Conventional model: increased brain activation during later remembered compared to forgotten scenes (**Table S3**). All results are shown at *p* < 0.05 FWE-corrected at cluster level (cluster-defining threshold of *p* < 0.001).

**Figure S3:**
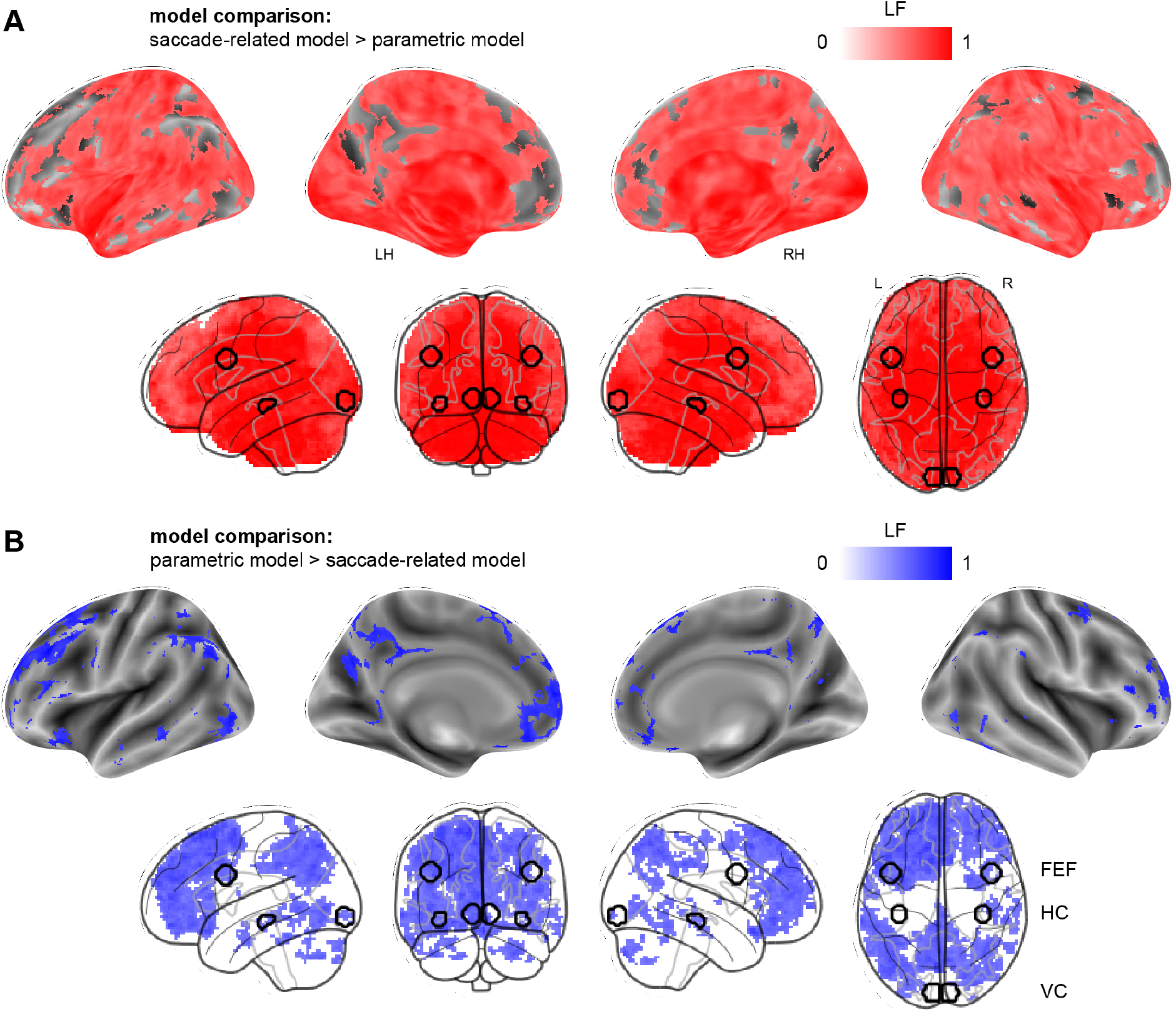
Results from model comparison. Model comparison using Bayesian Model Selection (BMS) between the **(A)** saccade-related and **(B)** parametric models yielded selected-model maps (SMMs) that indicated the model with the highest frequency (likeliest frequency, LF) per voxel (**Materials and Methods**). The saccade-related model (124868 voxels, marked in red) provided a better model fit compared to the parametric model (13194 voxels, marked in blue) in 90.4 % of all voxels in the brain (total voxel count: 138062 voxels). Across the respective voxels in which the saccade-related (red) and parametric model (blue) provided a better model fit, the LF for the saccade-related model was generally higher (average LF: saccade-related model, 0.71; parametric model: 0.55). We also extracted the average LF values per region-of-interest (bilateral ROIs marked with black contours on glass brains; frontal eye fields, FEF; hippocampus, HC; visual cortex, VC; **Results S1**).

**Figure S4:**
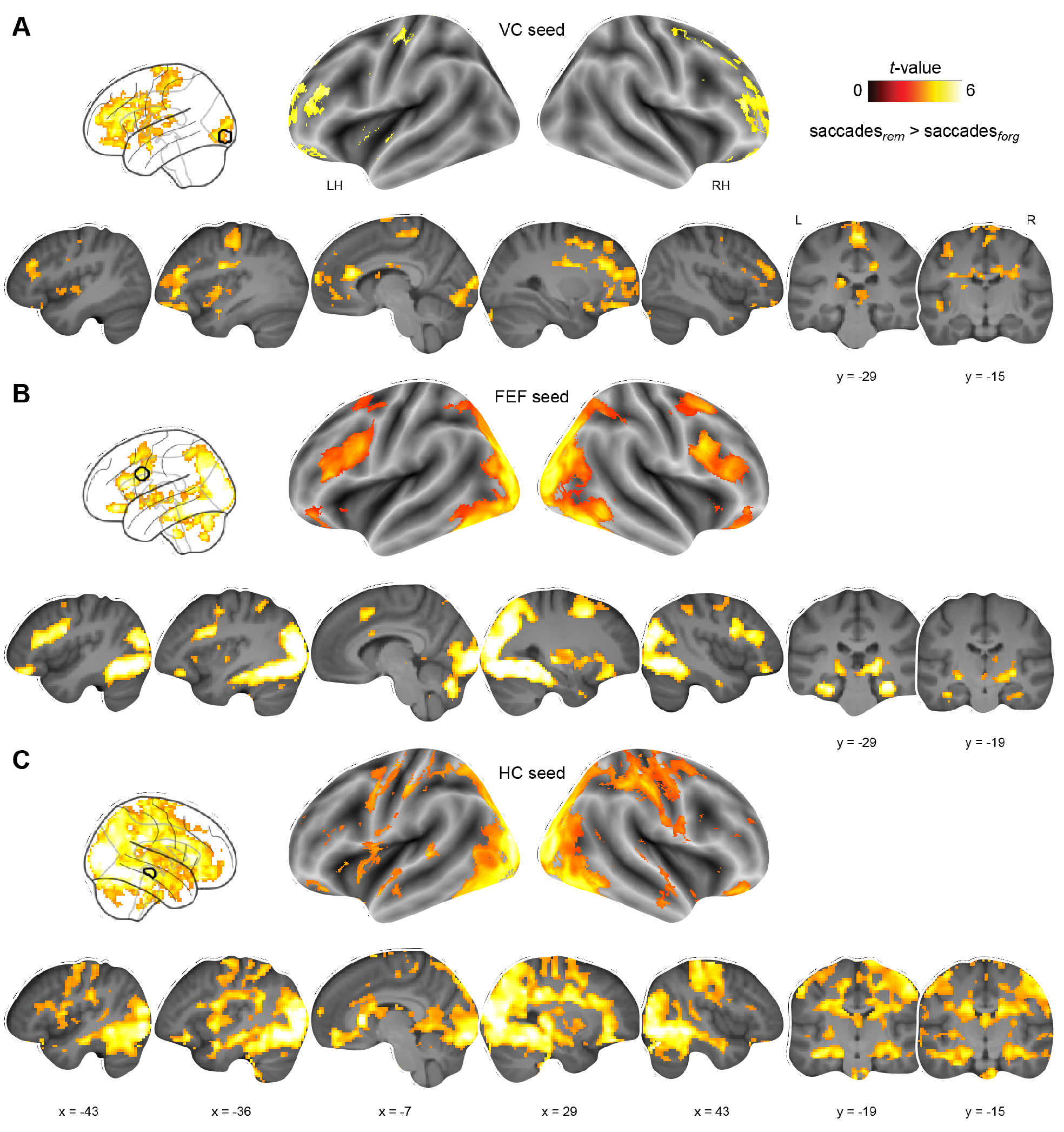
Whole-brain connectivity (PPI) during saccade-related memory formation from VC, FEF, and HC seeds. Connectivity increases during saccade-related memory formation (saccades_rem_ > saccades_forg_) based on seeds located in **(A)** the left visual cortex (VC), **(B)** the left frontal eye fields (FEF), and **(C)** the right hippocampus (HC; original seeds used for ROI-definition, see also **Materials and Methods**). Seeds are outlined in black, visible in glass brains. All results are shown at *p* < 0.05 FWE-corrected at cluster level (cluster-defining threshold of *p* < 0.001, see also **Table S5**).

**Figure S5:**
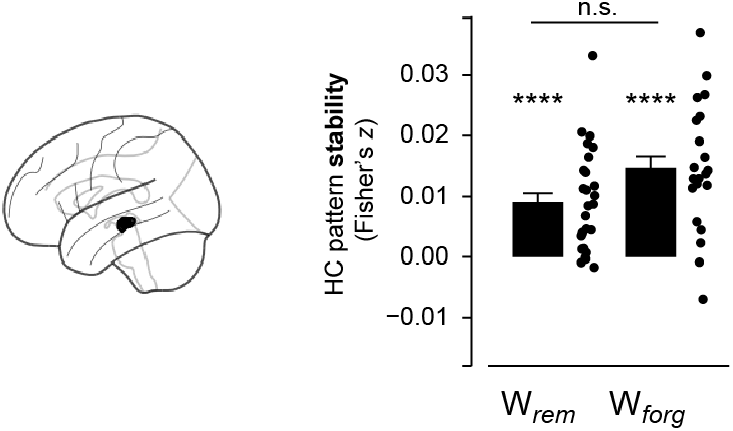
Saccade-related HC pattern stability. Average pattern stability extracted from the left hippocampus (HC) per memory condition (W_*rem*_, W_*forg*_); **** *p* < 0.0001, no significant difference (n.s.) between conditions.

**Figure S6:**
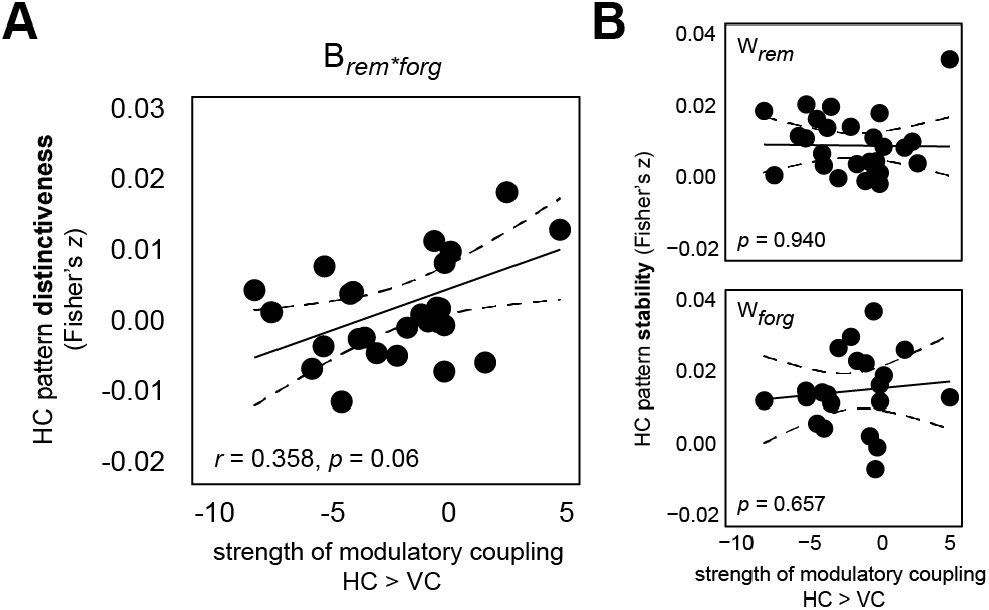
HC-to-VC inhibition is specifically related to distinct HC patterns during memory formation. Correlations between saccade-related inhibitory HC-to-VC coupling during successful memory formation (obtained from DCM analysis, left hemisphere) and saccade-related changes in hippocampal pattern similarity (left hemisphere, same ROI as also for DCM analysis). **(A)** HC-to-VC inhibition is specifically related to HC pattern distinctiveness (B_*rem*forg*_) rather than **(B)** pattern stability within each condition (W_*rem*_, W_*forg*_).

## Supplementary Tables

### General note regarding tables displaying fMRI results

Unless stated otherwise, significance for all MRI analyses was assessed using cluster-inference with a cluster-defining threshold of *p* < 0.001 and a cluster-probability of *p* < 0.05 family-wise error (FWE) corrected for multiple comparisons. The corrected cluster size (i.e., the spatial extent of a cluster that is required in order to be labeled as significant) was calculated using the SPM extension “CorrClusTh.m” and the Newton-Raphson search method (script provided by Thomas Nichols, University of Warwick, United Kingdom, and Marko Wilke, University of Tübingen, Germany)^1^.

MNI coordinates represent the location of cluster peak voxels. We report the first local maximum within each cluster. The complete results tables available at the Open Science Framework (https://osf.io/4dshu/).

Anatomical nomenclature for all tables was obtained from the Laboratory for Neuro Imaging (LONI) Brain Atlas (LBPA40, http://www.loni.usc.edu/atlases/; Shattuck et al., 2008). L, left; R, right; LH, left hemisphere; RH, right hemisphere.

**Table S1:** Saccade-related brain activation. Analysis consisted of separate one-sample *t*-tests (all *N* = 32), contrasts: [saccades_rem_ ∩ saccades_forg_] > baseline (critical cluster size: 116 voxels), saccades_rem_ > saccades_forg_ (107 voxels).

| Contrast & brain region | MNI |  |  | Z-value | Cluster size |
| --- | --- | --- | --- | --- | --- |
|  | x | y | z |  |  |
| ***[Saccades<sub>rem</sub> ∩ saccades<sub>forg</sub>] > baseline:*** |  |  |  |  |  |
| L middle occipital gyrus | -22 | -96 | 10 | 6.96 | 17173 |
| Cerebellum | 19 | -38 | -46 | 5 | 170 |
| R inferior frontal gyrus | 58 | 34 | 19 | 4.77 | 165 |
| L middle frontal gyrus | -14 | 72 | 7 | 4.39 | 133 |
| ***Saccades<sub>rem</sub> > saccades<sub>forg</sub>:*** |  |  |  |  |  |
| R middle occipital gyrus | 36 | -77 | 31 | 6.94 | 8514 |
| L middle occipital gyrus | -29 | -74 | 31 | 6.84 | 8026 |
| R precentral gyrus | 43 | 7 | 29 | 5.52 | 958 |
| R inferior frontal gyrus | 58 | 36 | 12 | 5.07 | 185 |
| Cerebellum | -7 | -74 | -38 | 5 | 224 |
| L postcentral gyrus | -58 | -12 | 48 | 4.62 | 570 |
| R precentral gyrus | 12 | -24 | 70 | 4.34 | 372 |
| L inferior frontal gyrus | -55 | 31 | 22 | 4.2 | 122 |
| Cerebellum | -14 | -48 | -50 | 4.09 | 108 |

**Table S2:**
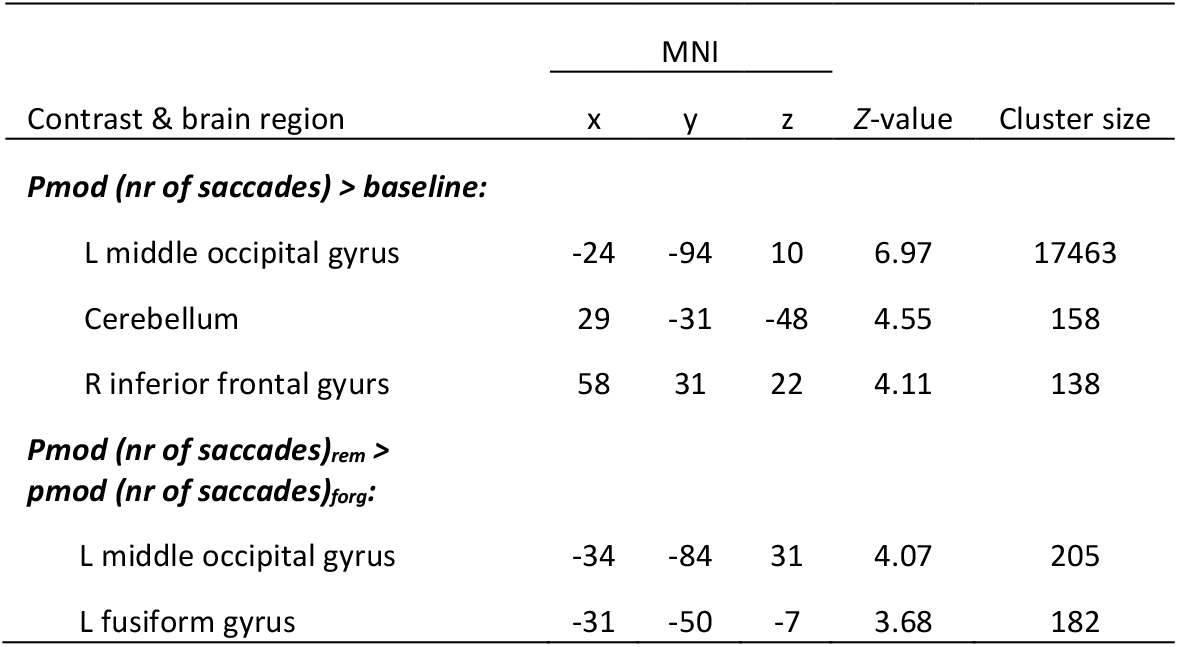
Brain activation using the parametric model. Analysis consisted of separate one-sample *t*-tests (all *N* = 32), contrasts: pmod (nr of saccades) > baseline (critical cluster size: 113 voxels), pmod (nr of saccades)_rem_ > pmod (nr of saccades)_forg_ (107 voxels).

| Contrast & brain region | MNI |  |  | Z-value | Cluster size |
| --- | --- | --- | --- | --- | --- |
|  | x | y | z |  |  |
| *Pmod (nr of saccades) > baseline:* |  |  |  |  |  |
| L middle occipital gyrus | -24 | -94 | 10 | 6.97 | 17463 |
| Cerebellum | 29 | -31 | -48 | 4.55 | 158 |
| R inferior frontal gyurs | 58 | 31 | 22 | 4.11 | 138 |
| *Pmod (nr of saccades)<sub>rem</sub> > pmod (nr of saccades)<sub>forg</sub>:* |  |  |  |  |  |
| L middle occipital gyrus | -34 | -84 | 31 | 4.07 | 205 |
| L fusiform gyrus | -31 | -50 | -7 | 3.68 | 182 |

**Table S3:** Brain activation using the conventional model. Analysis consisted of separate one-sample *t*-tests (all *N* = 32), contrast: rem > forg (critical cluster size: 111 voxels).

| Contrast & brain region | MNI |  |  | Z-value | Cluster size |
| --- | --- | --- | --- | --- | --- |
|  | x | y | z |  |  |
| ***Rem > forg:*** |  |  |  |  |  |
| R middle occipital gyrus | 38 | -79 | 29 | 7.33 | 19985 |
| R precentral gyrus | 43 | 7 | 29 | 5.15 | 387 |
| R inferior frontal gyrus | 58 | 36 | 12 | 5.15 | 159 |
| L postcentral gyrus | -55 | -14 | 53 | 4.59 | 237 |

**Table S4:**
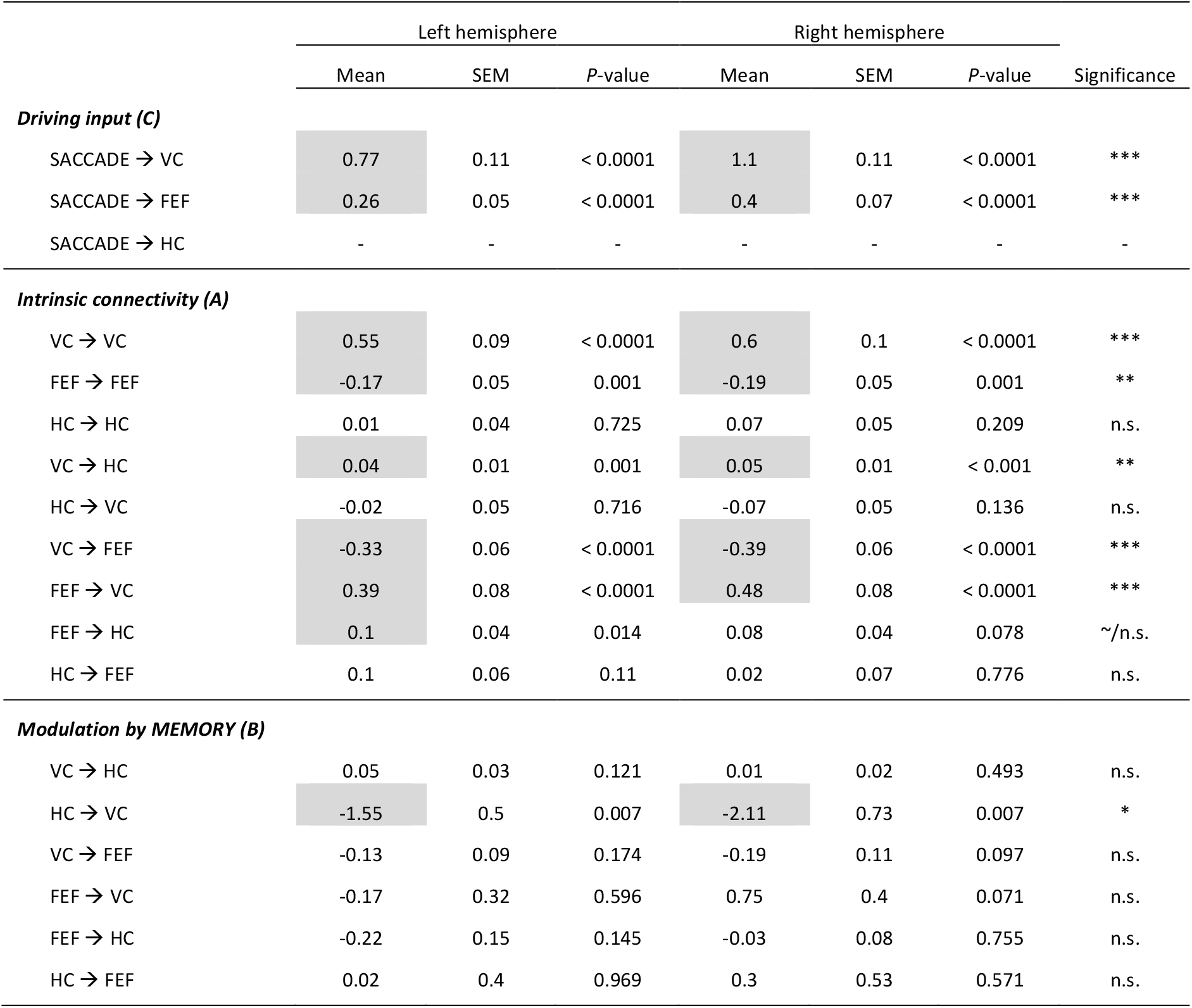
BMA results for left- and right-lateralized DCM networks. Results from Bayesian Model Averaging (BMA) based on left- and right-lateralized DCM networks (VC, visual cortex; FEF, frontal eye fields; HC, hippocampus). The strength of the coupling parameters for driving inputs (matric *C*), intrinsic connections (matrix *A*) and modulatory connections (matrix *B*; remembered > forgotten) is represented as mean (± SEM) values across all participants (*N* = 32). *P*-values were compared against the respective significance threshold (separate one-sample *t*-tests against zero, α_Bonferroni_; driving inputs: 0.05/2 comparisons = 0.025, intrinsic connectivity: 0.05/9 comparisons = 0.006, modulatory connectivity: 0.05/6 comparisons = 0.008. ***, *p* < 0.0001; **, *p* < 0.001; * *p* < 0.008; ∼, *p* < 0.05 but not surviving correction for multiple comparisons; n.s., not significant. To highlight the consistency of results between the left- and right-lateralized networks, significant findings are marked in grey.

|  | Left hemisphere |  |  | Right hemisphere |  |  |  |
| --- | --- | --- | --- | --- | --- | --- | --- |
|  | Mean | SEM | P-value | Mean | SEM | P-value | Significance |
| Driving input (C) |  |  |  |  |  |  |  |
| SACCADE → VC | 0.77 | 0.11 | < 0.0001 | 1.1 | 0.11 | < 0.0001 | *** |
| SACCADE → FEF | 0.26 | 0.05 | < 0.0001 | 0.4 | 0.07 | < 0.0001 | *** |
| SACCADE → HC | - | - | - | - | - | - | - |
| Intrinsic connectivity (A) |  |  |  |  |  |  |  |
| VC → VC | 0.55 | 0.09 | < 0.0001 | 0.6 | 0.1 | < 0.0001 | *** |
| FEF → FEF | -0.17 | 0.05 | 0.001 | -0.19 | 0.05 | 0.001 | ** |
| HC → HC | 0.01 | 0.04 | 0.725 | 0.07 | 0.05 | 0.209 | n.s. |
| VC → HC | 0.04 | 0.01 | 0.001 | 0.05 | 0.01 | < 0.001 | ** |
| HC → VC | -0.02 | 0.05 | 0.716 | -0.07 | 0.05 | 0.136 | n.s. |
| VC → FEF | -0.33 | 0.06 | < 0.0001 | -0.39 | 0.06 | < 0.0001 | *** |
| FEF → VC | 0.39 | 0.08 | < 0.0001 | 0.48 | 0.08 | < 0.0001 | *** |
| FEF → HC | 0.1 | 0.04 | 0.014 | 0.08 | 0.04 | 0.078 | ~/n.s. |
| HC → FEF | 0.1 | 0.06 | 0.11 | 0.02 | 0.07 | 0.776 | n.s. |
| Modulation by MEMORY (B) |  |  |  |  |  |  |  |
| VC → HC | 0.05 | 0.03 | 0.121 | 0.01 | 0.02 | 0.493 | n.s. |
| HC → VC | -1.55 | 0.5 | 0.007 | -2.11 | 0.73 | 0.007 | * |
| VC → FEF | -0.13 | 0.09 | 0.174 | -0.19 | 0.11 | 0.097 | n.s. |
| FEF → VC | -0.17 | 0.32 | 0.596 | 0.75 | 0.4 | 0.071 | n.s. |
| FEF → HC | -0.22 | 0.15 | 0.145 | -0.03 | 0.08 | 0.755 | n.s. |
| HC → FEF | 0.02 | 0.4 | 0.969 | 0.3 | 0.53 | 0.571 | n.s. |

**Table S5:** Whole-brain connectivity (PPI) Analysis consisted of a separate one-sample *t*-tests (each *N* = 32), contrast: saccades_rem_ > saccades_forg_ (critical cluster size: VC seed, 76 voxels; FEF seed, 65, voxels; HC seed, 83 voxels).

| Contrast & brain region | MNI |  |  | Z-value | Cluster size |
| --- | --- | --- | --- | --- | --- |
|  | x | y | z |  |  |
| **Visual cortex (VC) seed** |  |  |  |  |  |
| R middle frontal gyrus | 34 | 31 | 17 | 5.01 | 7529 |
| R precentral gyrus | 2 | -29 | 62 | 4.53 | 764 |
| R inferior temporal gyrus | 38 | 5 | -34 | 4.27 | 151 |
| L middle occipital gyrus | -14 | -91 | 2 | 4.12 | 749 |
| L precentral gyrus | -31 | -19 | 48 | 3.91 | 204 |
| Cerebellum | 26 | -91 | -26 | 3.77 | 93 |
| Thalamus | 7 | -24 | 2 | 3.75 | 76 |
| **Frontal eye fields (FEF) seed** |  |  |  |  |  |
| L inferior occipital gyrus | -36 | -74 | -10 | 6.65 | 16953 |
| R inferior frontal gyrus | 36 | 7 | 29 | 5.07 | 1100 |
| L lateral orbitofrontal gyrus | -41 | 38 | -14 | 4.88 | 258 |
| Cerebellum | 0 | -48 | -36 | 4.83 | 72 |
| L inferior frontal gyrus | -50 | 26 | 17 | 4.83 | 1246 |
| Cerebellum | -19 | -34 | -48 | 4.81 | 116 |
| R middle frontal gyrus | 29 | 12 | 48 | 4.72 | 604 |
| L insular cortex | -34 | 5 | 10 | 4.25 | 228 |
| Cerebellum | -26 | -70 | -48 | 4.24 | 85 |
| R superior frontal gyrus | 7 | 22 | 46 | 4.19 | 443 |
| L cingulate gyrus | -2 | 5 | 26 | 4.14 | 71 |
| R inferior temporal gyrus | 41 | -7 | -31 | 4.04 | 96 |
| L insular cortex | -26 | 19 | 0 | 3.9 | 65 |
| **Hippocampus (HC) seed** |  |  |  |  |  |
| R middle occipital gyrus | 24 | -86 | 12 | 5.98 | 41902 |
| L superior frontal gyrus | 0 | -2 | 74 | 4.22 | 96 |
| Brainstem | 7 | -17 | -36 | 4.04 | 182 |

**Table S6:** Saccade-related pattern similarity (RSA) Analysis consisted of a separate one-sample *t*-tests (all *N* = 32), contrast: W_rem_ > B_rem*forg_ (critical cluster size: 86 voxels).

| Contrast & brain region | MNI |  |  | Z-value | Cluster size |
| --- | --- | --- | --- | --- | --- |
|  | x | y | z |  |  |
| ***W<sub>rem</sub> > B<sub>rem</sub>\*forg:*** |  |  |  |  |  |
| L insular cortex | -31 | 14 | -12 | 4.63 | 851 |
| R inferior frontal gyrus | 43 | 22 | 19 | 4.55 | 423 |
| R superior occipital gyrus | 29 | -72 | 38 | 4.38 | 273 |
| L superior frontal gyrus | -14 | 14 | 48 | 4.31 | 272 |
| L putamen | -31 | -19 | -7 | 4.2 | 184 |
| Cerebellum | 41 | -67 | -22 | 3.9 | 126 |

1 http://www2.warwick.ac.uk/fac/sci/statistics/staff/academic-research/nichols/scripts/spm/

